# Acto-myosin driven functional nanoclusters of GPI-anchored proteins are generated by integrin receptor signaling

**DOI:** 10.1101/232223

**Authors:** Joseph Mathew Kalappurakkal, Anupama Ambika Anilkumar, Chandrima Patra, Thomas S. van Zanten, Michael P. Sheetz, Satyajit Mayor

## Abstract

GPI-anchored protein (GPI-AP) nanoclusters are generated by cortical acto-myosin activity. While our understanding of the physical principles behind this process is emerging, the molecular machinery required for the generation of these nanoclusters is unknown. Here, we show that ligand-mediated membrane receptor signaling triggers nanocluster formation. Both soluble and surface-tethered RGD ligands bind the β1-integrin receptor and activate focal adhesion and src-kinases, resulting in RhoA signaling. This cascade ultimately triggers actin-nucleation via specific formins, driving nanoclustering of both GPI-APs and a model transmembrane protein with an actin-binding domain. Integrin signaling concurrently results in talin mediated activation of vinculin. This is necessary for the coupling of the dynamic actin machinery to the inner leaflet driving GPI-AP nanoclustering. Disruption of GPI-AP nanoclustering in either GPI-anchor remodeling mutants or in cells that express vinculin mutants, provide evidence that these nanoclusters are necessary for activating cell spreading, a hallmark of integrin function.

## INTRODUCTION

Sub-compartmentalization of the plasma membrane (PM) via the lateral segregation of proteins and lipids into structural and signaling platforms is likely to play pivotal roles in the spatio-temporal regulation of many signaling systems. Finely tuned signaling systems such as T-cell receptor triggering at the immunological synapse (Gaus et al., 2005), B-cell receptor activation (Gupta and DeFranco, 2007; Mattila et al., 2013), and cell-ECM adhesion (Gaus et al., 2006; Lingwood and Simons, 2010; Simons and Toomre, 2000; van Zanten and Mayor, 2015), involve the generation of membrane domains. Such membrane domains, enriched in cholesterol, sphingolipids and outer leaflet lipid-tethered glycosylphosphatidylinositol (GPI)-anchored proteins, have often been termed as membrane ‘rafts’ (Sezgin et al., 2017).

The mechanism whereby cells generate these domains remains controversial. The size, scale and statistics of membrane heterogeneities in the cell are very different than what is predicted from thermodynamically driven phase segregation observed in artificial membranes or cell-free membrane preparations such as Giant Plasma Membrane Vesicles (GUVs) (Chiantia and London, 2012; Sezgin et al., 2012a, 2012b). Many of the ‘raft’ components such as outer leaflet GPI-APs or inner leaflet Ras molecules form nanoclusters (Plowman et al., 2005; Varma and Mayor, 1998). These nanoscale clusters are hierarchically organized into larger scale optically resolvable (mesoscale) domains where a significant fraction of the lipid-anchored proteins are present as nanoclusters (Goswami et al., 2008; Tian et al., 2007; van Zanten et al., 2009).

In the ‘resting’ state, GPI-APs at the cell surface are distributed as monomers with a small fraction of nanoclusters (20-40%) that is independent of total protein expression levels (Sharma et al., 2004; van Zanten et al., 2009). Under conditions of activation such as the binding of ligand to the integrin receptor, LFA-1, in immune cells, the fraction of GPI-APs in nanoclusters increases (~ 80%), and appears in spatial proximity to LFA-1 nanoclusters, regions designated as "hot-spots". Reduction of cholesterol levels, a treatment that also prevents the formation of GPI-AP nanoclusters, drastically inhibited the ligand binding capacity of these adhesion receptors (van Zanten et al., 2009). At the same time ligand-induced or crosslinking antibody-induced clustering of GPI-APs is sufficient to drive downstream signaling responses in the cell (Harder et al., 1998; Stefanová et al., 1991; Suzuki et al., 2007). The regulation of nanoscale clustering is, thus, likely to be an important determinant in high-fidelity signal transduction processes that operate at the cell surface (Harding and Hancock, 2008; Tian et al., 2007).

Extensive studies on the organization and dynamics of GPI-APs have indicated a crucial role for a dynamic actin layer at the cortex juxtaposed to the inner leaflet of the PM in the formation of such nanoclusters at the outer leaflet (Goswami et al., 2008; Saha et al., 2015). In previous work, we had proposed that dynamic actin filaments along with myosin motors form transient remodeling contractile platforms (asters) at the inner leaflet (Gowrishankar et al., 2012). These ‘asters’ immobilize clusters of the long-acyl chain containing phosphatidylserine (PS) at the inner leaflet which interact with long-acyl chain containing GPI-APs at the outer leaflet via a transbilayer coupling interaction, thereby creating nanoclusters (Raghupathy et al., 2015). These and other observations (Köster and Mayor, 2016; Rao and Mayor, 2014) indicate that the organization at the membrane might be better understood as an active actin-membrane composite, wherein the constituents in the fluid membrane bilayer interact with the dynamic actin cortex. In this context, membrane components can be classified into three classes based on their ability to couple with and regulate this active machinery: inert, passive and active (Gowrishankar et al., 2012). Inert molecules are those that are unable to interact with the underlying actin (for example unsaturated lipids at the outer leaflet or membrane proteins that lack any linkage to actin filaments), passive molecules are those that bind (and unbind) actin filaments (such as the GPI-APs as well as transmembrane proteins that possess actin-binding motifs at their cytoplasmic tails), and active molecules which not only bind but also influence the actin cytoskeleton dynamics at the membrane and in doing so could regulate local membrane organization.

Despite the emergence of a theoretical understanding of the active mechanics behind the generation of the nanoscale assemblies and their distribution and dynamics, the molecular machinery for their formation has been missing; the nucleators of actin, triggers of myosin function, and the linkage between the actin and the PS lipid are uncharacterized. In this manuscript we uncover the molecular machinery that governs the formation of the dynamic actin-based membrane patterning system.

Here we show that integrin receptors behave as a prime example of an ‘active’ molecule that regulates actin nucleators and myosin activity necessary to build the hierarchical organization of clusters. We find that upon engagement with RGD-containing ligands, integrin receptors through their ability to activate the FAK and src kinases and the resultant RhoA activation trigger formins necessary for the generation of the dynamic actin filaments. RhoA also activates the ROCK pathway, required for myosin activation. Importantly, we also identify vinculin, a ubiquitous protein that associates with focal adhesions, as a molecule necessary for directing the generated dynamic actin filaments to the inner-leaflet lipids and thereby generating GPI-AP-nanoclusters. Furthermore, using GPI-anchor remodeling mutants as well as vinculin mutants, which fail to support nanocluster formation, we show that the nanoclusters created by this active machinery are necessary for activating cell spreading, a hallmark of integrin function.

## RESULTS

### Integrin activation generates nanoclusters of the outer leaflet GPI-APs

The integrin family of heterodimeric transmembrane receptors binds various extracellular ligands that activate a multitude of structural and signaling molecules (Hynes, 2002; Vicente-Manzanares et al., 2009). Integrins exhibit the hallmarks of an ‘active’ molecule that upon ligand engagement could alter the nanoscale organization of cell surface molecules in its vicinity through its ability to regulate the cortical acto-myosin network. Earlier studies of the integrin LFA1 activation in fixed immune cells have shown that upon binding to its ligand, ICAM-1, hot spots of GPI-AP clusters are formed, localized to the site of integrin activation within 30 mins of activation (van Zanten et al., 2009). This prompted us to test whether the activation of other integrins had a similar effect on the nanoclustering of GPI-APs albeit in a different cellular context. We used fluorescence emission anisotropy based microscopy to assess the extent of homo-FRET between fluorescently-tagged GPI-APs (Ghosh et al., 2012). Homo-FRET results in the lowering of anisotropy, providing a facile way to monitor nano scale clustering in intact living cells (Sharma et al., 2004; Varma and Mayor, 1998). Cells (CHO) stably expressing mEGFP or mYFP-tagged GPI (MYG-1) were deadhered and re-plated under serum-free conditions either on glass coated with 1%BSA or on glass coated with the extracellular matrix (ECM) protein fibronectin (FN) (**Figure 1A**). FN is capable of engaging with a specific subset of integrins (Humphries et al., 2006; Hynes, 2002) that promotes cell spreading (Figure1C), whereas the BSA surface is relatively inert to cell spreading at a similar time point (Figure 1C).

**Figure 1.**
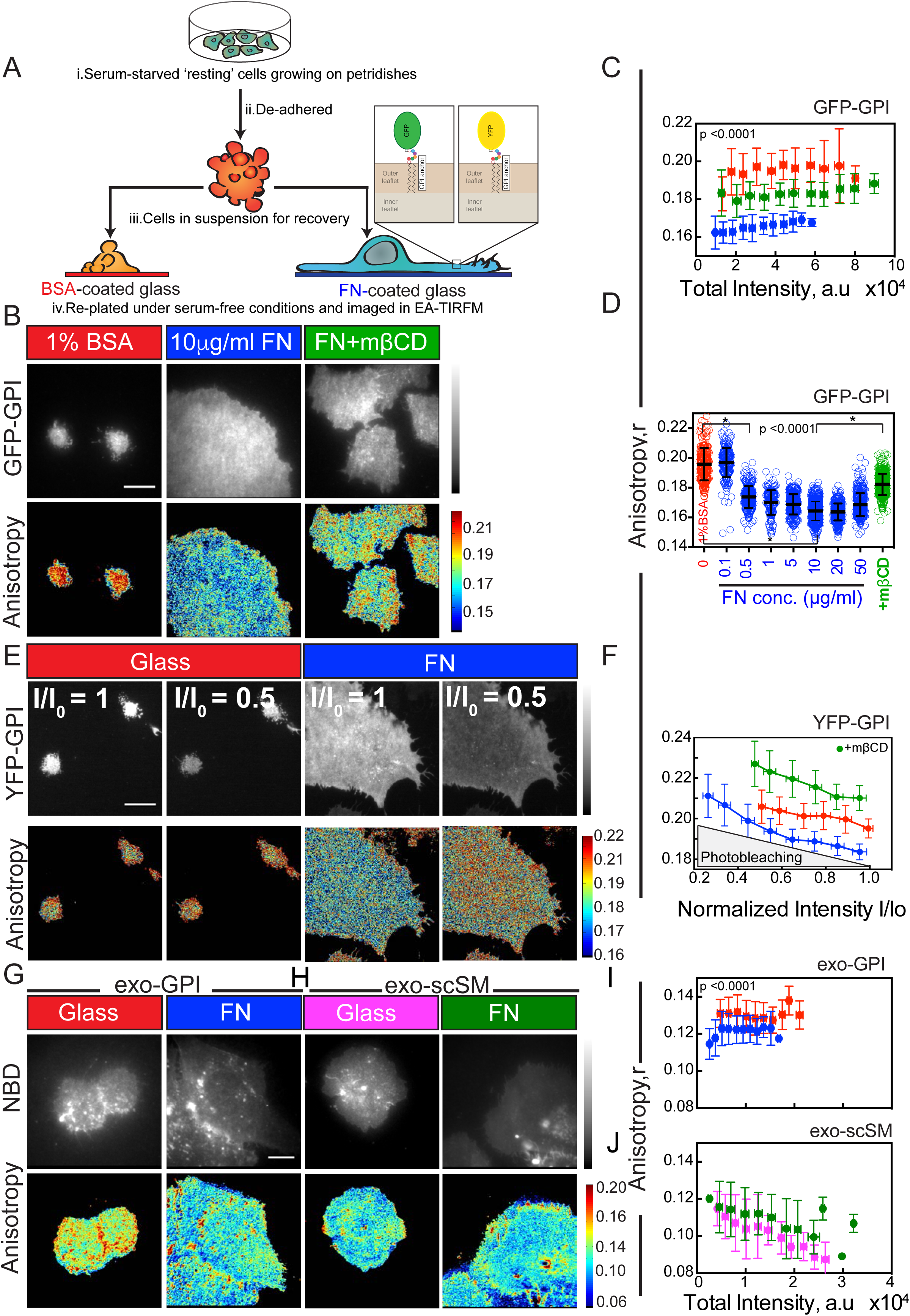
Activation of fibronectin binding integrins leads to enhanced nanoclustering of GPI-APs. (A) Cartoon illustrates the assay used in this study. Serum-starved (3-8 hrs) fibroblastic cells were de-adhered and re-plated back under serum-free conditions either on Fibronectin (FN)–coated or 1%BSA coated glass-bottom coverslips. The extent of nanoclustering of fluorescent protein-tagged GPI-APs expressed in these cells was monitored by measuring the fluorescence anisotropy using an Emission Anisotropy TIRF microscope (EA-TIRFM). Inset is the schematic representation of GFP or YFP tagged GPI-APs localized to the outer-leaflet of the plasma membrane.(B-C) Intensity and steady state anisotropy images (B) and anisotropy versus intensity plot (C) of GFP-GPI in live cells plated on 1%BSA (red) or on 10μg/ml FN (blue) or plated on FN and subsequently treated with 10mM of the cholesterol depleting agent methyl-β-cyclodextrin (mβCD) (green) and collected using EA-TIRFM. (D) Box plot representing the mean anisotropy values of GFP-GPI in CHO cells, plated on glass-bottom coverslips coated with 1% BSA (red) or FN (blue) at the indicated concentrations before (blue) or after (green) treatment with mβCD. (E-F) Intensity and steady state anisotropy images (E) at the initial time point (I/Io =1) and at 50% photobleaching (I/Io=0.5) and graph demonstrating the change in the fluorescence anisotropy upon photo-bleaching of YFP-GPI (F) expressed in CHO cells plated on glass (red) or on 10μg/ml FN (blue) or plated on FN and subsequently treated with mβCD (green), plotted against the fluorescence intensity value (*I*) normalized to its value before photobleaching (*I_0_*). Note that the starting anisotropy value of YFP-GPI in cells plated on glass was higher than that on FN. The anisotropy increased upon photobleaching to attain a final anisotropy value corresponding the value obtained in cells after cholesterol depletion (green horizontal bar). This value likely represents the anisotropy value of YFP-GPI-monomers (A∞). (G-J) Intensity and steady state anisotropy images (G, H) and anisotropy versus intensity plot (I, J) of CHO-K1 cells labeled for 3 hours with NBD-GPI (I; GPI analogue, exoGPI) or C6NBD-SM (J; short chain SM; scSM), de-adhered and plated for 1 h at 37°C on glass (red) or 10μg/ml FN (blue) and imaged on EA-TIRFM. Note the fluorescence anisotropy of C6-NBD-SM does not change, whereas exogenously incorporated NBD-GPI exhibits a lower anisotropy when cells are plated on FN compared to glass alone. Scale bar 10 μm. Error bars represent the SD. See also Figure S1

Although the amount of EGFP-GPI expressed on the cell surface is comparable (**Figure1B-C**; Total Intensity axis), the anisotropy (**Figure 1B-C**; Anisotropy axis) is much lower in cells plated on FN compared to that measured on BSA coated glass. The low (or high) anisotropy is discerned by ‘blue (or red)’ pixels in the heat map-encoded anisotropy image in **Figure 1B** and used throughout this manuscript.

The observed decrease in anisotropy occurs in a FN-concentration dependent manner saturating at ~10μg/ml FN solution concentration (**Figure 1D**). This decrease is specific for FN since when plated on 0.01% Poly-L-Lysine (that permits integrin-independent adhesion; (Schlaepfer et al., 1994) or on Laminin, Collagen-1 or Vitronectin (that engages a different subset of integrins), there is no significant reduction in the anisotropy of GFP-GPI (**Figure S1A, B**).

An increase in anisotropy upon photo-bleaching would indicate homo-FRET as a cause for the lowering of anisotropy (Sharma et al., 2004). Photo-bleaching YFP-GPI results in a net increase in anisotropy (**Figure 1E, F**) confirming our expectation. The typical profile of a linear increase is consistent with the presence of nanoclusters of YFP-GPI when cells are plated on FN (Sharma et al., 2004). The higher value of initial anisotropy and the minimal change in the YFP-GPI anisotropy value in cells plated on glass upon photo-bleaching, corroborates the low fraction of nanoclusters that are formed under this condition (**Figure1E, F**). Additionally, the decrease in anisotropy observed when cells are plated on FN is sensitive to the removal of cholesterol by the cholesterol-sequestering agent methyl β-cyclodextrin that disrupts nanoscale organization of GPI-APs [(Raghupathy et al., 2015); FN+mβCD; **Figure 1B, C**], confirming the enhancement in nanoclustering of GPI-APs on this substrate. The decrease in anisotropy occurs only for specific membrane constituents; there is a decrease in anisotropy of an exogenously incorporated fluorescent GPI analogue (NBD-GPI) (**Figure 1G, I**; exo-GPI) whereas an ‘inert’ fluorescent short chain-containing sphingomyelin analogue (C_6_-NBD-SM) incorporated into cells plated on FN does not exhibit this decrease (**Figure 1H, J**; exo-scSM).

The αV-class (αVβ3) and β1-class (α5β1) integrins are the primary integrins that mediate fibroblast cell spreading on FN (Humphries et al., 2006; Leiss et al., 2008).We utilized various function perturbing antibodies targeted against either the β1 or the αV class of integrins to discern which of these integrin sub-types are involved in the FN mediated generation of GPI-AP nanoclusters in human U2OS cells (Byron et al., 2009). We observed a loss in nanoclustering of GPI-APs (increase in GFP-GPI anisotropy) when U2OS cells were pre-treated with the increasing concentrations of β1-blocking antibody and subsequently plated on FN (**Figure S1D-E**). There was also a significant decrease in the cell spread area as a function of β1-blocking antibody concentration indicating that U2OS cells predominantly utilize the β1 integrin to spread on FN. At the highest concentration, the anisotropy values obtained were comparable to those obtained after treatment with mβCD (**Figure S1D**). There was no increase in GPI-AP anisotropy when U2OS cells were treated with antibodies that do not block spreading [(neutral non-function perturbing β1 antibody (K20) or Transferrin-receptor antibody (OKT9); **Figure S1E-F**) or αV-blocking antibody (17E6; data not shown)] and subsequently plated on FN. Additionally, U2OS cells plated on the αVβ3 ligand vitronectin (Charo et al., 1990) does not exhibit an increase in GPI-AP nanoclustering. Together, these data indicate that the enhanced nanoclustering occurs when cells are plated on FN, and this is mediated by the activated β1-class of integrins.

We next probed if an increase in affinity of the integrin for its ligand can alter the nanoclustering of GPI-APs. To test this, we plated U2OS cells on low concentrations of ligand (0.5μg/ml) in the presence of increasing amounts of Mn^2+^, an ion that potentiates integrin activation (Dransfield et al., 1992; Mould, 2002; Takagi et al., 2002). Prior treatment of cells with increasing amounts of Mn^2+^ resulted in a dose-dependent decrease in anisotropy of GPI-APs in the presence of low FN (**Figure S1G-H**). On high FN (10μg/ml; and higher), addition of Mn^2+^ did not result in a further decrease in anisotropy (**Figure S1G-H**) indicating no further increase in GPI-AP nanoclustering.

Taken together, these data indicate that shifting the equilibrium towards a ligand-engaged integrin, either by increasing FN density or by activation through Mn^2+^ promotes the generation of GPI-AP nanoclusters.

### Localized nanoclustering in the vicinity of the activated integrin receptor

When plated on FN, the cells go through three major phases of behavior from a round state in suspension to a fully-flattened circular morphology (Dubin-Thaler et al., 2004). These are: Phase 0 (P0), the wetting phase mediated by the initial engagement of the integrin with its ligand; Phase 1 (P1), the rapid expansion phase where sensing of the mechanical rigidity and chemical suitability of the substrate and the establishment of a large contact area takes place (Giannone et al., 2004); and finally, Phase 2 (P2), where myosin II contractility based probing of the substrate via periodic protrusion/retraction of the cell edge and continued asymptotic spreading to maximum area is attained. These phases define critical checkpoints for progression from a suspended state to a fully spread state. This is organized in a specific spatial and temporal order involving distinct sets of protein modules in each phase (Wolfenson et al., 2015), allowing us to correlate the changes in nanoclustering of GPI-APs with these universal characteristics of a spreading cell.

We examined the effect of cell spreading (CHO) on fibronectin-coated surfaces on GPI-AP nanoclustering (**Figure 2A-C; Supplementary Movie 1**). At the level of a single cell, the cell surface that first comes in contact with the FN-coated area appears to be devoid of nanoclusters (red areas in **Figure 2A**) and starts to acquire nanoclusters (‘blue’ pixels at the cell periphery) co-incidental with the P0-P1 transition phase in cell spreading (**Figure 2B and C;** Pink-yellow transition zone**, Figure S2B**). Analysis of this behavior over a large number of cells shows that there is a sudden and consistent decrease in the steady-state anisotropy of GFP-GPI (δAnisotropy) that precedes the peak in cell expansion (δArea; that occurs in P2 phase) by ~100-200 seconds (**Figure 2C**).

**Figure 2.**
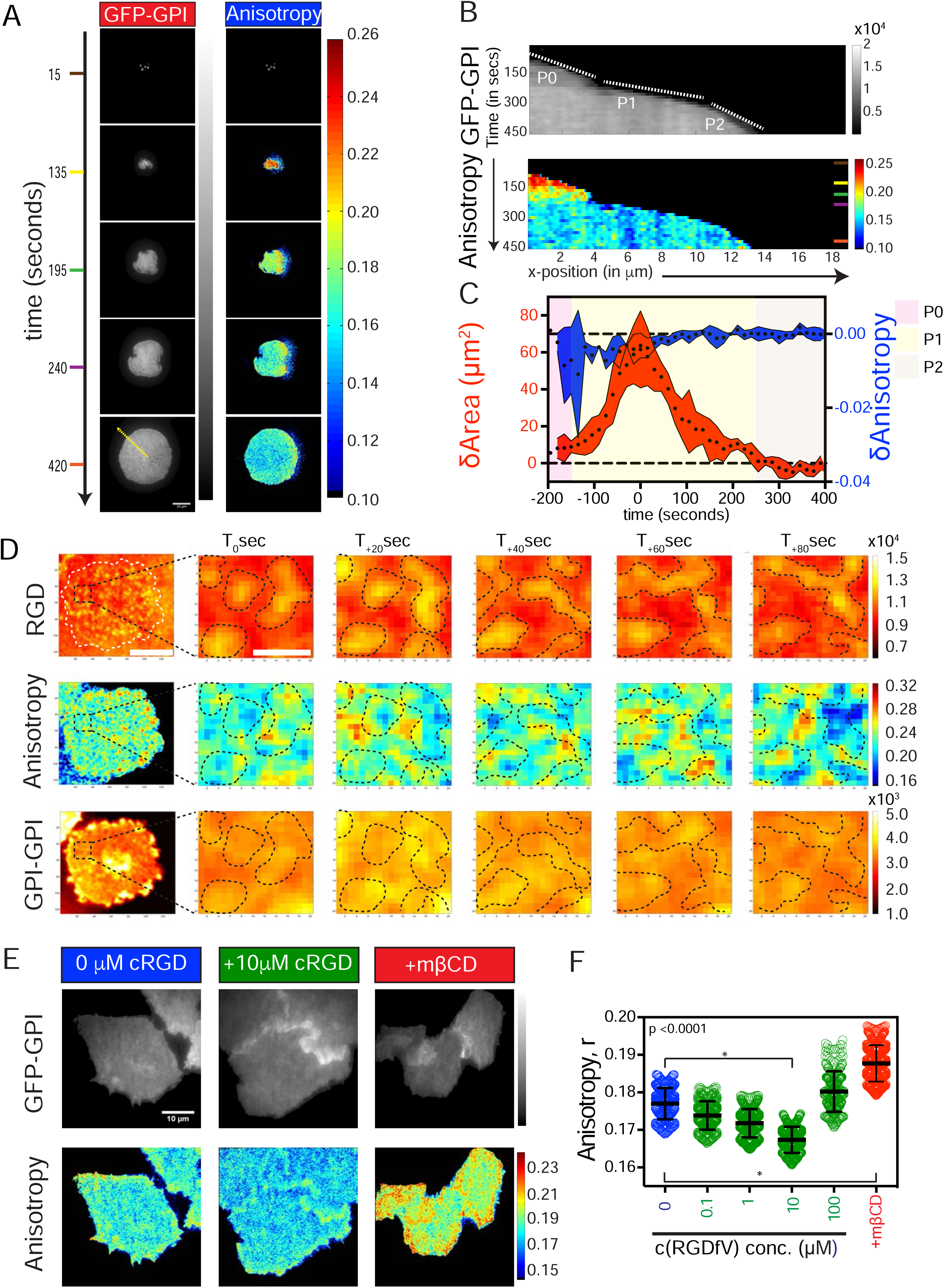
Activation of RGD binding integrins leads to enhanced nanoclustering of GPI-APs in its local vicinity. (A-C) CHO cells stably expressing GFP-GPI were treated as described in Figure 1A and re-plated on FN (10μg/ml)-coated glass-bottom cover slips, to observe the temporal evolution of cell spreading and nanoclustering of GFP-GPI as monitored by EA-TIRFM. Images (A) show snapshots of GFP-GPI Intensity and steady state Anisotropy at the indicated times, post settling on FN. Kymograph (3 pixel average) (B) of intensity (top) and anisotropy (bottom) of a line drawn perpendicular to the cell edge (yellow line in A) showing the three phases of cell spreading and correlative changes in the anisotropy of GFP-GPI. Note that the rapid decrease in anisotropy (increase in the nanoclustering) occurs at the transition between P0-P1 phase. Graph (C) shows the change in area (6Area;red curve) correlated to changes in GFP-GPI anisotropy (6Anisotropy;blue curve) as a function of time. Note that the peak change in fluorescence anisotropy (blue curve) occurs about 100-200 seconds before the peak increase in cell spread area. Data depicts mean change in whole cell anisotropy values and cell spread area between two consecutive frames (15s) of 11 cells measured at the indicated spreading time; Data has been aligned relative to the timing of the peak area change. (D,E) Representative snapshot taken from region of interest (2 μm × 2 μm) over time of CHO cells expressing GFP-GPI plated (right panel) on RGD-functionalized SLB taken sequentially, as described in Figure S2C. Note the correlation between RGD cluster intensity as reported with DyLight 650 labeled neutravidin (RGD; top panel; see also Figure S 2G) and the GFP-GPI steady state anisotropy (Anisotropy; middle panel) imaged sequentially in an Emission anisotropy –equipped spinning disc confocal microscope; GFP-GPI intensity (bottom panel), shows no correlated patterns. Scale bar, 5 μm for whole cell image and 1 μm for ROI. (E-F) Intensity and steady state anisotropy images (E) and box plots with mean anisotropy (F) of GFP-GPI expressing CHO cells plated on Glass for 2-days without (0 μM cRGD; blue) or with indicated concentrations of soluble RGDfV (green) for 30 mins at 37 °C, or with 10mM mβCD for 45 min at 37 °C and imaged using EA-TIRFM. Higher concentrations of cRGD (>100 μM) induced cell rounding and detachment. Correspondingly the cells also exhibited a higher anisotropy value. Scale bar 10 μm. Error bars represent SD. See also Figure S2.

In the P0 phase, integrin engagement and clustering is an early step in the formation of cell-ECM adhesions and is independent of force (Choi et al., 2008). To probe if the effects of integrin activation on the promotion of GPI-AP nanoclustering are force-dependent and localized to the sites of integrin activation, we employed a supported lipid bilayer (SLB) system functionalized with a mobile lipid-attached cyclic-RGD ligand (**Figure S2C**); Arg-Gly-Asp (RGD) ligand is the sequence motif in FN that mediates integrin-engagement (Ruoslahti, 1996). Here, the transiently immobilized fluorescently-tagged ligands serve as reporters of the ligated integrins (Yu et al., 2011, **Figure S2C**). This system facilitates the observation of local membrane organization during the early stages of integrin mediated cell adhesion by enabling the simultaneous tracking of the dynamics of the nascent integrin clusters and the nanoclustering of GFP-GPI in the membrane in the vicinity of this cluster. (**Figure 2D; Figure S2H**). Although the engaged integrin is unable to exert significant traction on the fluid bilayer which results in the inability of cells to fully spread and arrest in the P0 phase (**Figure S2F**), there is enhanced nanoclustering of GPI-APs on cells engaged with lipid-attached mobile RGD ligands compared to cells plated on glass (**Figure S2D, E**). In many cases we observe a characteristic pattern where integrin cluster formation often precedes the local decrease in anisotropy of GFP-GPI (**Figure 2D**; see more examples in **Figure S2G, H**). These results show that the activation of the FN binding integrin receptors triggers a localized change in GPI-AP nanoclustering.

The time from the initial contact until initiation of cell spreading is inversely correlated with the ligand density (Dubin-Thaler et al., 2004), suggesting that the process is triggered by the integration of chemical signals via integrin receptor engagement to its ligand. To test the possibility that the extent of GPI-AP nanoclustering is also an integral response of a chemical signaling process, we treated cells with a soluble cRGDfV peptide that has been shown to activate signaling molecules downstream of integrin receptor binding (Huveneers et al., 2008; Zhang et al., 2014). Strikingly, we also observe an increase in nanoclustering of GFP-GPI in cells plated on glass and treated with the soluble cRGDfV in a cholesterol and dose-dependent manner (**Figure 2E, F**) indicating that the increase in nanoclustering is triggered by a signaling response initiated by integrin-RGD-binding.

### GPI-AP nanoclustering is mediated via a RhoA signaling pathway downstream of integrin activation

RhoGTPases, tyrosine kinases, and various bona fide cytoskeletal modifying proteins are involved in the steps of integrin-mediated cell spreading behavior (Vicente-Manzanares et al., 2009). To investigate the signaling pathway activated by integrins that leads to GPI-APs nanoclustering, we employed a chemical and genetic perturbation approach. Pre-treatment of cells with the src-family kinase (SFK) inhibitor, PP2 (Hanke et al., 1996) and the focal adhesion kinase (FAK) inhibitor PF 573 228 (Slack-Davis et al., 2007) results in a dramatic decrease in GPI-AP nanoclustering in cells plated on FN (**Figure 3A,B**). Correspondingly, FAK null fibroblasts also fail to support FN-induced GPI-AP nanoclustering (**Figure S3C, D**).

**Figure 3.**
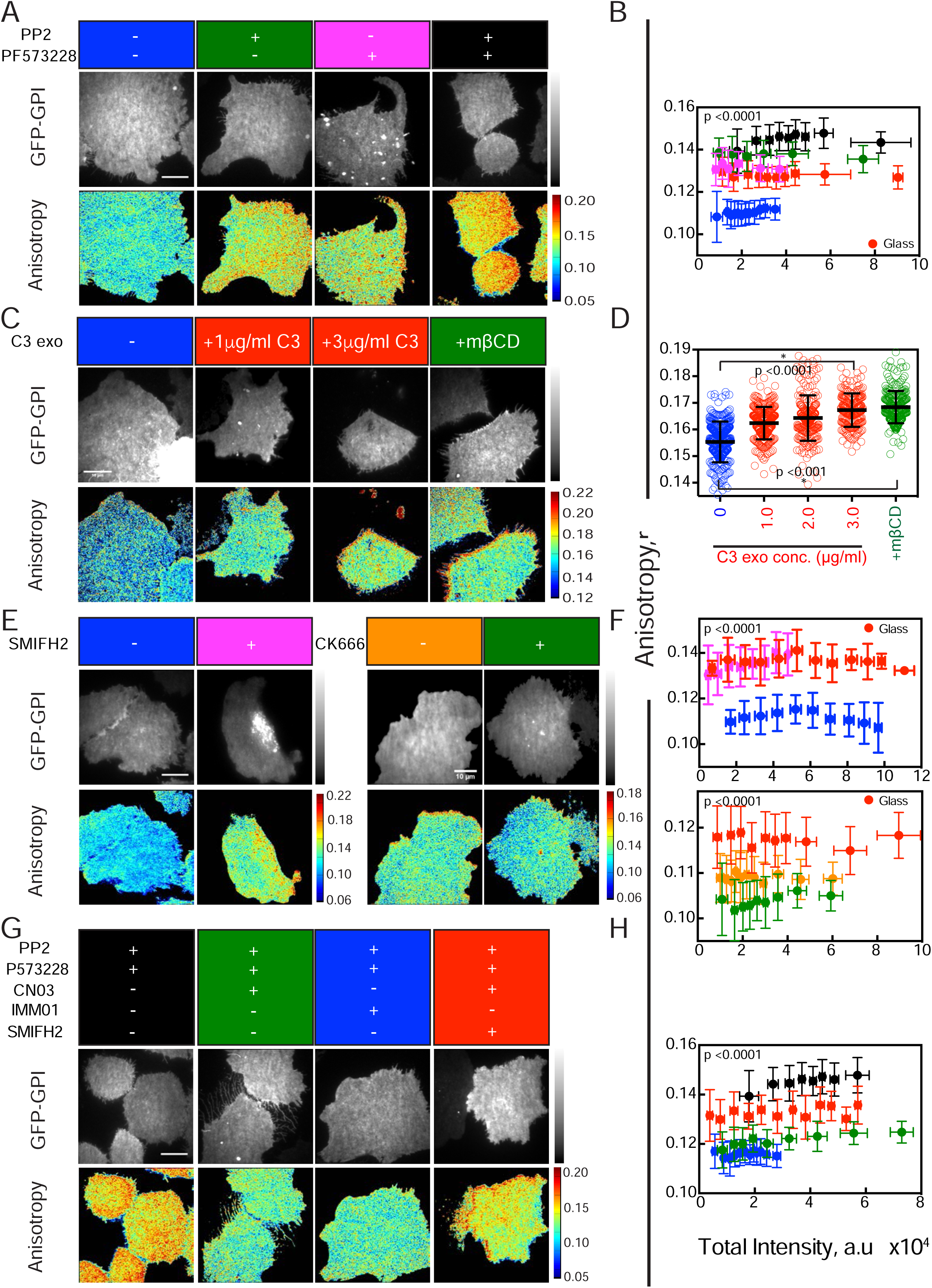
Inhibition of SFK/FAK, Rho GTPase and formins leads to loss of FN-triggered nanoclustering of GPI-APs. (A-F) Intensity and steady state anisotropy images (A, C, E) and intensity versus anisotropy plots (B, D, F) of GFP-GPI expressing CHO cells plated on FN-coated or non-coated (Red; Glass) glass-bottom dishes imaged via EA-TIRFM. The emission anisotropy of GFP-GPI was determined in cells pretreated with 20 μM src family tyrosine kinase inhibitor PP2 (+PP2; green in A-B), 10 μM focal adhesion kinase inhibitor PF-573-228 (+PF-573228; magenta in A-B) or both (+PP2+PF 573 228; black in A-B), pretreatment (2 h) with indicated concentrations of the cell-permeable Rho inhibitor exoenzyme C3 transferase (+C3; red in C-D), 10μM formin inhibitor (+SMIFH2;magenta in E-F), or 10μM formin agonist (+IMM01; green in G-H). Note in all cases, treatment with inhibitors increased the emission anisotropy of GFP-GPI consistent with a failure of FN-engagement to enhance nanoclustering, whereas activation of formin via IMM01 agonist, decreases emission anisotropy of GFP-GPI on cells plated on glass (green in G-H), in the absence of FN. (G, H) Intensity and steady state anisotropy images (G) and plot of anisotropy versus intensity (H) of GFP-GPI expressing CHO cells pre-treated and plated on 10μg/ml FN in presence of SFK-FAK inhibitor (20μM PP2+10μM PF-573 228) without (black in G-H) or with 10μg/ml RhoA activator (+CN03; green in G-H) or with 10μM formin agonist (+IMM01; blue in G-H), or with RhoA activator CN03 and 10μM formin inhibitor (+SMIFH2;red in G-H) and imaged via EA-TIRFM. Note that the RhoA or formin activators reverse the effects of SFK-FAK inhibition. The RhoA activator reversal remains sensitive to inhibition via the formin-inhibitor (SMIFH2) indicating formins operate downstream of RhoA activation. Scale bar 10 μm. Error bars represent SD. See also Figure S3.

A downstream target of the SFK and FAK kinases during integrin mediated signaling is the Rho family GTPase member, RhoA (Cox et al., 2001; Guilluy et al., 2011; Ren et al., 1999). Increasing concentrations of the cell permeable Rho inhibitor C3 exoenzyme which specifically inhibit RhoA activity (Aktories et al., 1987; Braun et al., 1989)), also inhibited GPI-AP nanoclustering in a dose dependent manner when cells were plated on FN (**Figure 3C, D**). In contrast, addition of a cell-permeable RhoA activator [CN03; (Flatau et al., 1997; Schmidt et al., 1997)] induced enhanced nanoclustering of GPI-APs even when cells were plated on plain glass, a condition where there is minimal integrin activation (**Figure S3E, F**). Furthermore, the failure in promoting nanoclustering upon FN engagement when the cells were treated with both SFK-FAK inhibitors can be rescued by the ectopic addition of CN03 (**Figure 3G, H**). These results indicate that RhoA operates downstream of SFK-FAK in the molecular pathway that mediates the nanoclustering of GPI-APs.

### Formin nucleators are necessary for GPI-AP nanoclustering

We next investigated the role of the actin-nucleators that are downstream targets of integrin activation in mediating the nanoclustering of GPI-APs. Pre-treatment of cells with SMIFH2, a small molecule inhibitor of the formin class of actin nucleators (Rizvi et al., 2009) led to loss of FN-mediated nanoclustering of GPI-APs (**Figure 3E, F**), whereas the Arp2/3 inhibitor CK666 (Nolen et al., 2009) had no effect on GPI-AP nanoclustering (**Figure 3E, F**). Moreover, the acute loss of GPI-AP nanoclusters observed when cells were treated with inhibitors of SFK and FAK could be rescued by treatment of cells with a formin activator (IMM01; Lash et al., 2013) which in turn is reversed by SMIFH2 treatment (**Figure 3G, H**), suggesting that formins are downstream of SFK/FAK-RhoA in this pathway that mediates nanoclustering of GPI-APs. In addition, treatment of cells plated on uncoated glass with the formin activator (IMM01) resulted in an increase in nanoclustering of GPI-APs even in the absence of integrin ligand engagement (**Figure S3G, H**). This suggests that formin activation is an important step in the integrin mediated signaling response that drives enhanced nanoclustering of GPI-APs.

To investigate the identity of the specific formin that mediates the nanoclustering of GPI-AP, we utilized specific RNAis to reduce the expression of two candidate formins, mDia1 (DIAPH1) and FHOD1 in U2OS cells. Upon reduction of the levels of mDia1 and FHOD1 between ~80 and ~40%, respectively (**Figure S4A, B**), there was a drastic decrease in nanoclustering of GPI-APs. The loss of nanoclustering was more significant when FHOD1 levels were reduced when compared to mDia1 (**Figure S4C, D**), implicating FHOD1 as one of the major formin members involved in this process. Taken together, these results indicate that actin filaments nucleated by specific formins are involved in the FN-integrin induced nanoclustering of GPI-APs.

### Integrin activation triggers an acto-myosin-based clustering mechanism

To test whether changes in nanoclustering of GPI-APs induced by integrin activation are mediated by the upregulation of the cortical acto-myosin based machinery described previously (Gowrishankar et al., 2012), we monitored the organization of a model chimeric receptor composed of an extracellular reporter domain derived from folate receptor [FR], a transmembrane segment [TM] and a cytosolic domain derived from the actin binding domain of ezrin [Ez-AFBD], FRTM-Ez-AFBD (**Figure 4A**). This chimeric protein also forms nanoscale clusters that are dependent on its ability to bind actin and associate with a dynamic acto-myosin machinery (Gowrishankar et al., 2012). Steady-state anisotropy measurements of fluorescently labeled FRTM-Ez-AFBD expressing cells showed a decrease in anisotropy when plated on FN (**Figure 4B, C**), similar to that observed for the GPI-APs (**Figure 1C**). By contrast, cells expressing a mutated version of the FRTM-Ez-AFBD that is incapable of binding actin (FRTM-Ez-AFBD*; **Figure 4D**) did not display changes in the steady-state anisotropy upon engagement with FN (**Figure 4E, F**).

**Figure 4.**
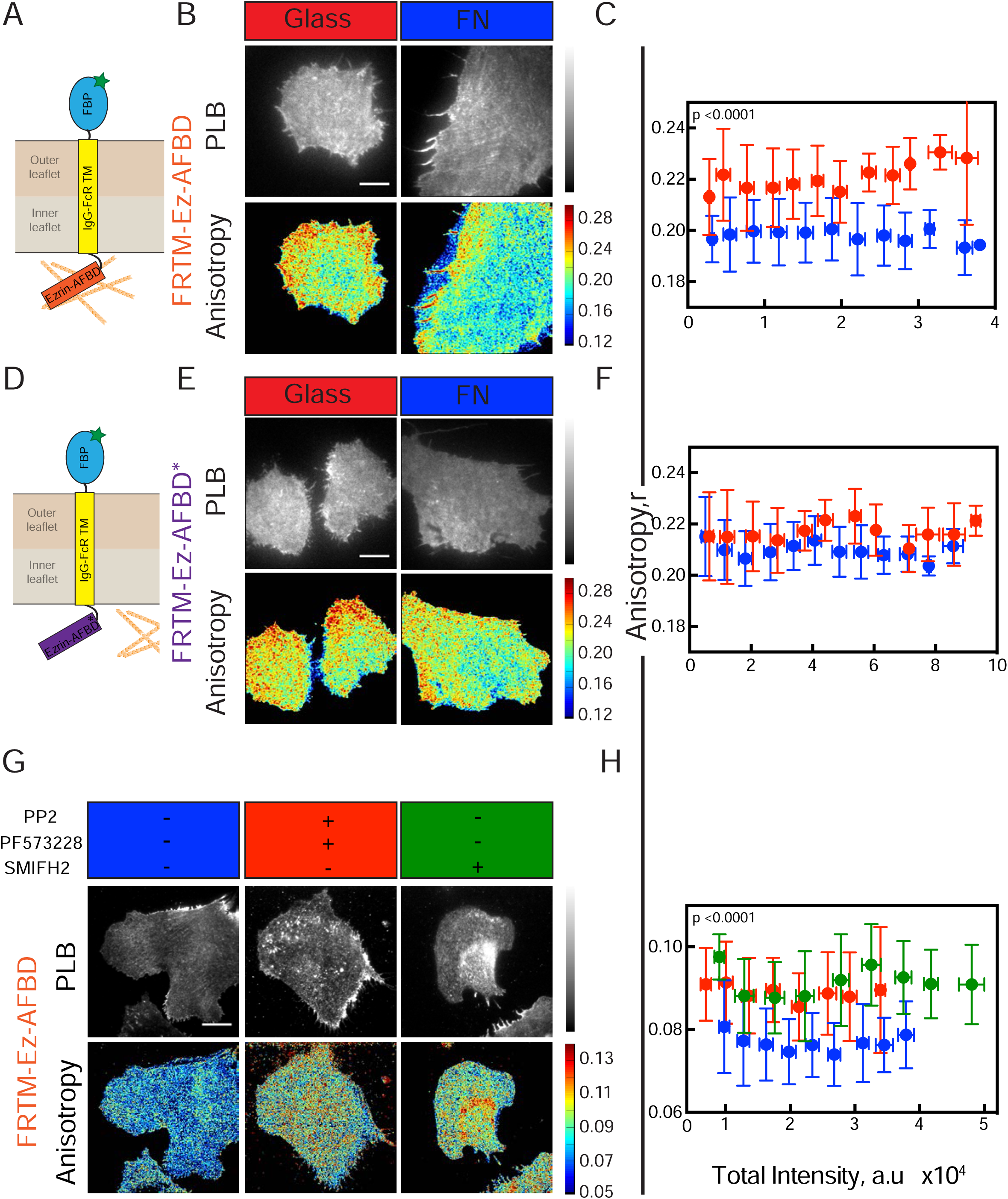
Integrin activation induces changes in dynamic actin activity at the cytoplasmic leaflet. (A) Schematic of model transmembrane protein with the extracellular region derived from the folate binding protein (FBP), transmembrane segment of IgG-Fc receptor (IgG-FcR TM) and an actin binding domain derived from the ezrin protein (FR-TM-Ez-AFBD) and (D) a mutated version of the same construct as in A, FR-TM-Ezrin-R579A (FR-TM-Ez-AFBD) that renders the Ez-AFBD domain incapable of interacting with actin. B-H) Intensity and steady state anisotropy images (B, E, G) and plots of anisotropy versus intensity (C, F, H) of CHO cells stably expressing FRTM-Ez-AFBD (B, C, G, H) or FRTM-Ez-AFBD* (E,F) labeled with fluorescent folic acid analogue, Pteroyl-lysyl-Bodipy™(PLB), for 3 hours at 37°C and re-plated either on FN-coated (blue in C, F, H) or uncoated (red in C, F) glass-bottom coverslip dishes and imaged via EA-TIRFM. In panel G and graph H, cells were re-plated after pre-treatment with only DMSO (blue in G-H), or 20μM PP2 and 10μM PF-573228 (red in G-H), or 10μM SMIFH2 (green in G-H). The difference in values of anisotropy of PLB-labelled FRTM-Ez-AFBD in G, compared to B is due to the use of a different EA-TIRFM imaging station, equipped with different NA optics. Error bars represent SD. See also Figure S4.

A signature of the dynamic actin-filaments at the membrane surface is the decreased diffusion of GFP-tagged-actin filament binding domain of Utrophin (GFP-Utr) as measured by fluorescence correlation spectroscopy (FCS) (Gowrishankar et al., 2012). When FCS traces were taken from regions in the periphery of the cell that were devoid of stress-fibers (**Figure S4E**), we detected at least two diffusing species (**Figure S4F**); one corresponding to the diffusion timescale of unbound GFP-Utr (0.3 ms < *τ* < 3 ms) and another slower component (<>10ms) that corresponds to GFP-Utr bound to actin filaments with an approximate filament length of ~200nm (Gowrishankar et al., 2012). Treatment of cells with the formin inhibitor SMIFH2, resulted in a loss of only the slow diffusing component (**Figure S4F, G**). This coincides with the loss of the dynamic pool of actin filaments that is likely to mediate the nanoclustering of membrane proteins (Saha et al., 2015). The nanoclustering of the chimeric transmembrane receptor (FRTM-Ez-AFBD) was also abrogated upon inhibition of formins as well as SFK/FAK on FN (**Figure 4G, H**).

Next we tested the role of integrin-stimulated ROCK activation in nanoclustering of GPI-APs. RhoA via ROCK can stimulate myosin light chain (MLC) phosphorylation directly or indirectly through the inhibition of MLC phosphatase. Inhibition of ROCK using the Y-27632 inhibitor (Uehata et al., 1997) results in loss of nanoclusters of GPI-APs when cells are plated on FN (**Figure S4H, I**) as does treatment with the MLC kinase (MLCK) inhibitor ML-7 (**Figure S4H, I**).

Together these data provide evidence that integrin signaling mediated by src and FAK kinases through active RhoA regulates actin polymerization via formins. This couples integrin ligation to the generation of nanoclusters. The role of myosin activity in promoting nanoclustering indicates that signaling activates a dynamic acto-myosin machinery to promote the nanoclustering of GPI-anchored and other actin-filament binding domain (AFBD) containing proteins.

### Talin and vinculin are necessary for the generation of GPI-AP nanoclusters

To further understand the mechanism of GPI-AP nanocluster generation, we tested the role of focal adhesion proteins, talin and vinculin, that form an integral part of the mechano-chemical signal transduction machinery downstream of integrin engagement (Vicente-Manzanares et al., 2009). Anisotropy measurements on vinculin knock out (vin-/-) mouse embryonic fibroblasts (MEFs) (Janoštiak et al., 2014) transfected with GFP-GPI, and freshly plated on FN showed a relatively high anisotropy value, which was unaffected by treatment with mβCD (**Figure 5A,B**). This indicates that cholesterol-sensitive nanoclusters do not form without vinculin. To restore vinculin function, we transiently transfected full-length mCherry-tagged vinculin (Thievessen et al., 2013a) into vin-/- MEFs re-plated on FN and measured anisotropy of co-transfected GFP-GPI. Re-introduction of vinculin restored cholesterol-dependent GPI-AP nanoclustering ascertained by decreased GFP-GPI anisotropy that was sensitive to cholesterol removal (**Figure 5A, B**). These data indicate that the loss of nanoclustering in vin-/-cells was only due to the absence of vinculin, providing a convenient test bed to explore the role of vinculin in GPI-AP nanoclustering.

**Figure 5.**
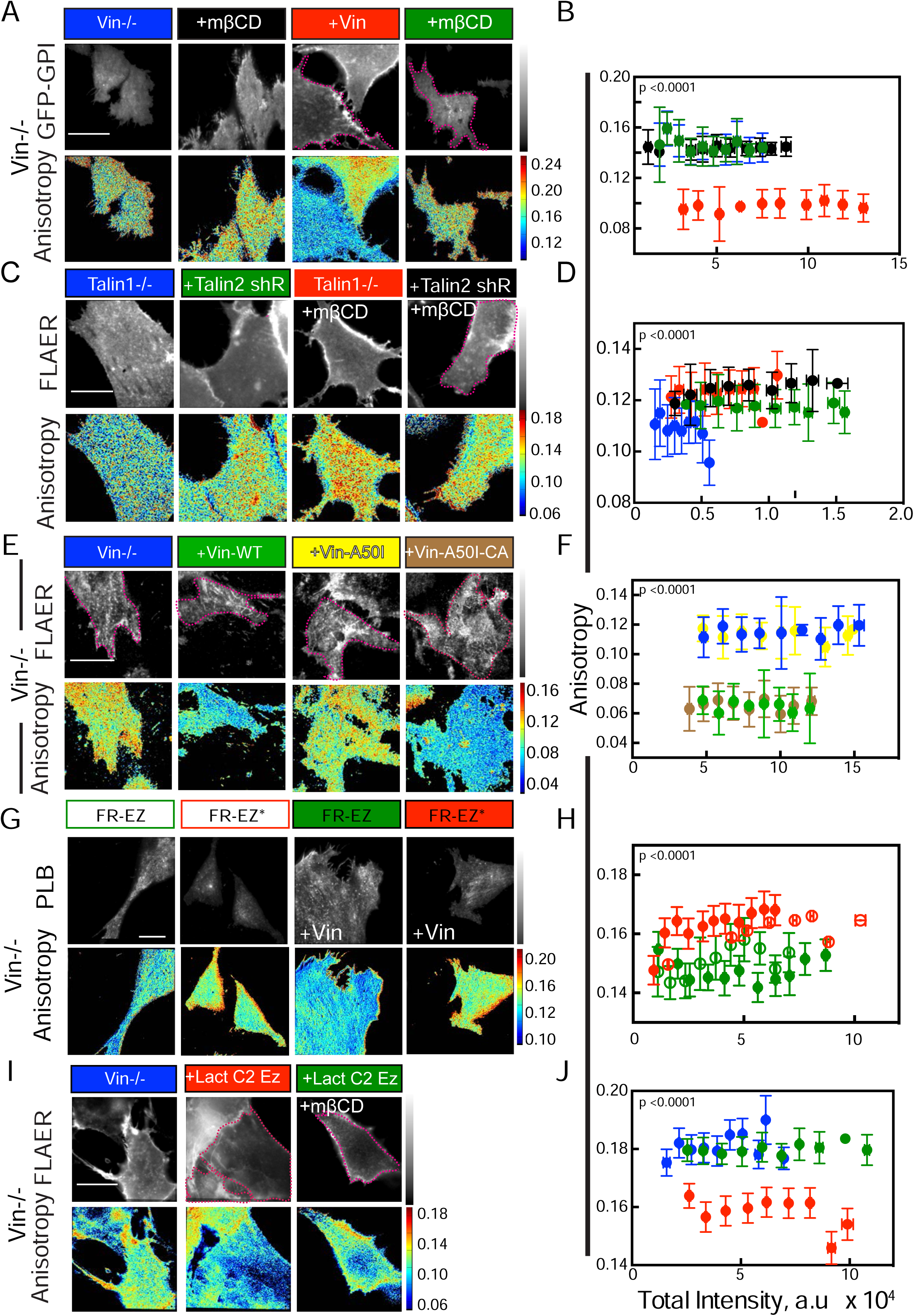
Talin and vinculin are required for facilitating GPI-AP nanoclustering in mouse embryonic fibroblasts (MEFs) (A-J) Intensity and anisotropy images of vinculin deficient (vin-/-; A, B, E-J) or talin 1 deficient (Talin1-/-) MEFs (C-D) either transfected with GFP-GPI (A, B) or labeled with Alexa-568-FLAER (FLAER; C-F, I-J) or PLB (pteroyl lysine conjugated to Bodipy-TMR; G, H) and imaged via EA-TIRFM after re-plating the cells on FN-coated glass-bottom dishes. Cells were also transfected with mcherry-vinculin (A, B), Talin2 shRNA (C, D), GFP-vinculin (Vin-WT; E, F),the indicated vinculin variants (E-F), FREZ, FREZ* (G, H) or Lact C2 Ez (I, J) 12-16 hours prior to being taken for imaging. mCherry vinculin (mCh-Vin; red in B) Talin2 shRNA (black in D) and Lact C2 Ez (green in J) expressing cells were additionally treated with mβCD. Transfected cells are marked by dotted magenta lines. Scale bar 10 μm. Error bars represent SD. See also Figure S5.

Vinculin exists in an auto-inhibited state in cells, and is activated by several interacting molecules, which bind to specific domains (**Figure S5C**). A well-characterized mode of activation of vinculin is through the binding of its head domain to talin, opening up the tail domains for interaction with actin and lipids (Case et al., 2015; Golji and Mofrad, 2010). We first addressed the role of talin in nanoclustering of GPI-APs, using talin1 knockout MEFs. Since loss of talin1 leads to over expression of the talin2 isoform (Zhang et al., 2008), we additionally depleted these cells of talin2 with talin2-shRNA co-expressed with GFP (**Figure S5A, B**). We monitored the nanoclustering of GPI-APs in these cells using a fluorescently-tagged oligomerization-defective aerolysin variant (A568-FLAER) previously characterized to report on the native distribution of endogenous GPI-APs (Raghupathy et al., 2015). A568-FLAER exhibited a higher fluorescence emission anisotropy in the talin2 shRNA expressing cells compared to the talin-1 alone deficient cells (**Figure 5C, D**), consistent with a loss of nanoclustering of GPI-anchored proteins in talin1-/- cells after talin2 depletion. However, this increase was less than that observed for the loss of vinculin; the partial loss of talin2 (as indicated by immunostaining for talin2) could serve as a confounding factor in these experiments (**Figure S5A, B**).

Expression of Vin-A50I, which is incapable of binding talin (Case et al., 2015; **Figure S5C**) in vin-/-MEFs, fails to support GPI-AP nanoclustering (**Figure 5E, F**). However, a constitutively activated vinculin, Vin-A50I-CA, which does not require talin for its activation (Case et al., 2015) restored GPI-AP nanoclustering in the vin-/- MEFs (**Figure 5E, F**). Together these results suggest that GPI-AP nanoclustering normally requires vinculin activation by talin.

### Vinculin activation specifically links integrin signaling and GPI-AP nanoclustering

We next asked if vinculin is necessary for triggering the actomyosin-based clustering machinery downstream of integrin activation. We examined the status of nanoclustering of FRTM-Ez-AFBD in vin-/- cells (**Figure 5G, H**). Monitoring anisotropy of FRTM-Ez-AFBD and the FRTM-Ez-AFBD* mutant constructs indicated that the nanoclustering mechanism is unaffected in the absence of vinculin; FRTM-Ez-AFB exhibited a lower anisotropy value compared to the FRTM-Ez-AFBD* in vin-/-cells as well as in the vinculin restored cells (**Figure 5G, H**). Furthermore, the restoration of GPI-AP nanoclustering observed in vin-/- MEFs by the expression of vinculin is completely disrupted upon pre-treatment with the src family inhibitor, PP2, or FAK inhibitor, PF 573228 or the formin inhibitor, SMIFH2 (**Figure 6A, B**). GPI-AP nanoclustering is not restored by treatment of cells with the formin activator, IMM01 in the vin-/- cells. By contrast, transmembrane protein clustering is brought about upon integrin activation in these cells, indicating that vin-/- cells are not defective in generating the acto-myosin machinery responsible for clustering. Addition of an artificial linker (LactC2-Ez-AFBD) (Raghupathy et al., 2015) is able to fully restore the nanoclustering of GPI-APs, in the vin-/- cells. This indicates that vinculin activation is not necessary for the creation of the acto-mysoin machinery but is rather involved in the pathway that links actin to the inner-leaflet lipids (**Figure 5I, J**).

**Figure 6.**
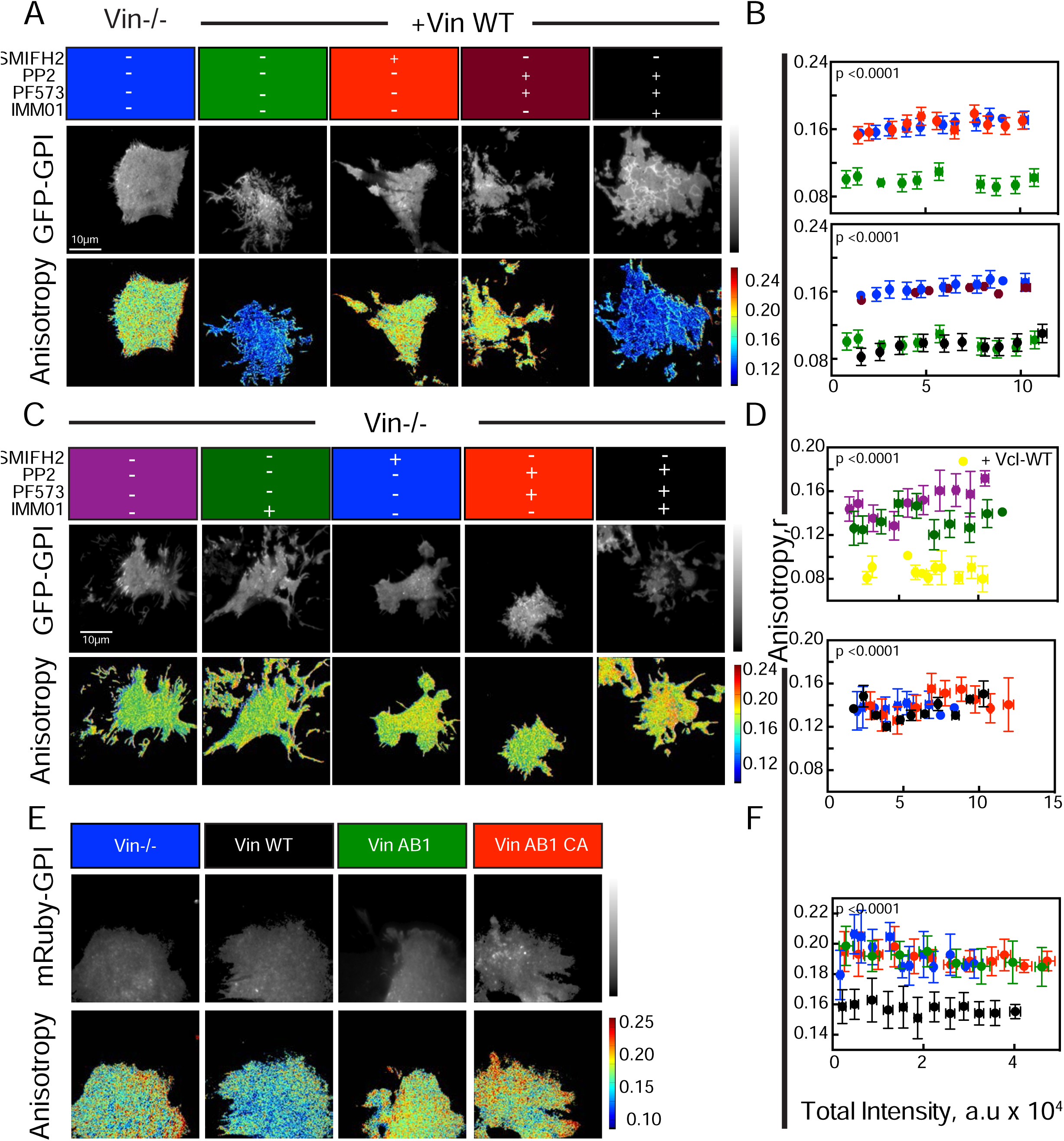
Vinculin facilitates GPI-AP nanoclustering in an integrin signaling sensitive manner. (A-D) Intensity and anisotropy images (A, C) and the intensity versus anisotropy plot (B, D) of GFP-GPI expressing MEFs in the presence (A,B) or absence (C, D) of vinculin treated with various inhibitors like SMIFH2, PP2, PF573228 and IMM01 that interferes with the integrin signaling pathway that generates dynamic actin. Note that an increase in anisotropy was observed when cells were treated with SMIFH2 (red (A, B) or PP2 and PF573228 in the presence (maroon; A, B) or absence of vinculin (red; C, D) as indicated and a decrease in anisotropy was observed on treatment with a cocktail of PP2, PF573228 and IMM01 (black) in the presence of vinculin (A, B), but an increase in anisotropy was observed on treating cells with a cocktail of PP2, PF573228 and IMM01 (black) in the absence of vinculin (C, D). (E-F) Intensity and anisotropy images (E) and the intensity versus anisotropy plot (F) of mRuby-GPI expressing vin-/- MEFs transiently transfected with Vin WT (black), Vin AB1 (green) or Vin AB1 CA (red). An increase in anisotropy was observed when cells were transfected with Vin AB1 and Vin AB1 CA compared to Vin WT suggesting a role for vinculin’s actin binding domain in GPI-AP nanoclustering. Error bar represent SD. See also Figure S6.

### Lipid and actin binding capacity of vinculin are necessary for GPI-AP nanoclustering

Vinculin possesses a negatively charged lipid binding site in its tail domain and mutation of this site results in a loss in its ability to bind to negatively charged lipids in the membrane (Humphries et al., 2007). Importantly, expression of this Vin-Ld mutant that lacks lipid-binding capacity (**Figure S5C)** in vin-/- MEFs failed to restore GPI-AP nanoclustering (**Figure S6A, B**). To determine if the failure to bind lipids, keeps vinculin in an inactive state, we generated Vin-Ld-CA*, that is constitutively activated (Humphries et al., 2007) (**Figure S5C**). This mutant also failed to restore GPI-AP nanoclustering in vin-/- MEFs (**Figure S6A, B)**. These results indicate that the lipid binding capacity of vinculin is necessary for it to catalyze GPI-AP nanocluster formation and accounts for the mechanistic differences between the nanoclustering of GPI-APs and those of transmembrane proteins with actin binding motifs.

To assess if the actin-binding capacity of vinculin is necessary to bring about GPI-AP nanoclustering, we expressed a mutant version of vinculin which has reduced capacity to bind to actin (Case et al., 2015) Vin AB1 (**Figure S5C**). Even though this mutant of vinculin localizes to focal adhesions, it failed to rescue nanoclustering of GPI-APs consistent with the role of the actin binding domain of vinculin in GPI-AP nanoclustering (**Figure 6E, F**).

### Cells defective in the GPI-AP nanoclusters formation exhibit aberrant integrin function

Cells that lack vinculin have defective integrin mediated responses; they lack the P1 phase of integrin mediated spreading and exhibit aberrant FAs (**Figure S7 H-I;** see also (Thievessen et al., 2013)). We hypothesized that some of these defects may be a consequence of the inability of cells to build functional nanoclusters. To test this, we utilized mutant cells that are deficient in two enzymes (PGAP2 and PGAP3) required for the remodeling of the unsaturated GPI-anchor acyl chains to long, saturated chains. This defect results in the inability of PGAP2/PGAP3 mutant cells to make GPI-AP nanoclusters (Raghupathy et al., 2015), and inefficient GPI-AP incorporation into detergent-resistant membranes (Maeda et al., 2007).

When freshly plated on fibronectin, these mutant cells failed to exhibit a decrease in the anisotropy of GFP-GPI and this defect could be reversed by restoring the activities of the PGAP2 and PGAP3 enzymes (**Figure S7A, B**). In comparison with either wild type cells or mutant cells rescued with wild type copies of PGAP2 and PGAP3, the PGAP2/PGAP3 mutant cells lack the P1 spreading phase when plated on substrates coated with fibronectin (**Figure 6A, B**). These mutant cells also lack a protrusive lamellipodia and possess fewer smaller adhesions (**Figure S7F-G**) and exhibit bleb-based cell spreading (**Supplementary Movie 2**). They also do not exhibit a rapid increase in cell area when spreading on FN, characteristic of the P1 phase (**Figure 7C; Supplementary Movie 3**). This defect is not due to defects in integrin activation in the mutant cells, since antibodies that bind to either active or in-active conformations of the β1-integrins bind equivalently to the mutant and wild type cells (**Figure S7C**).

**Figure 7.**
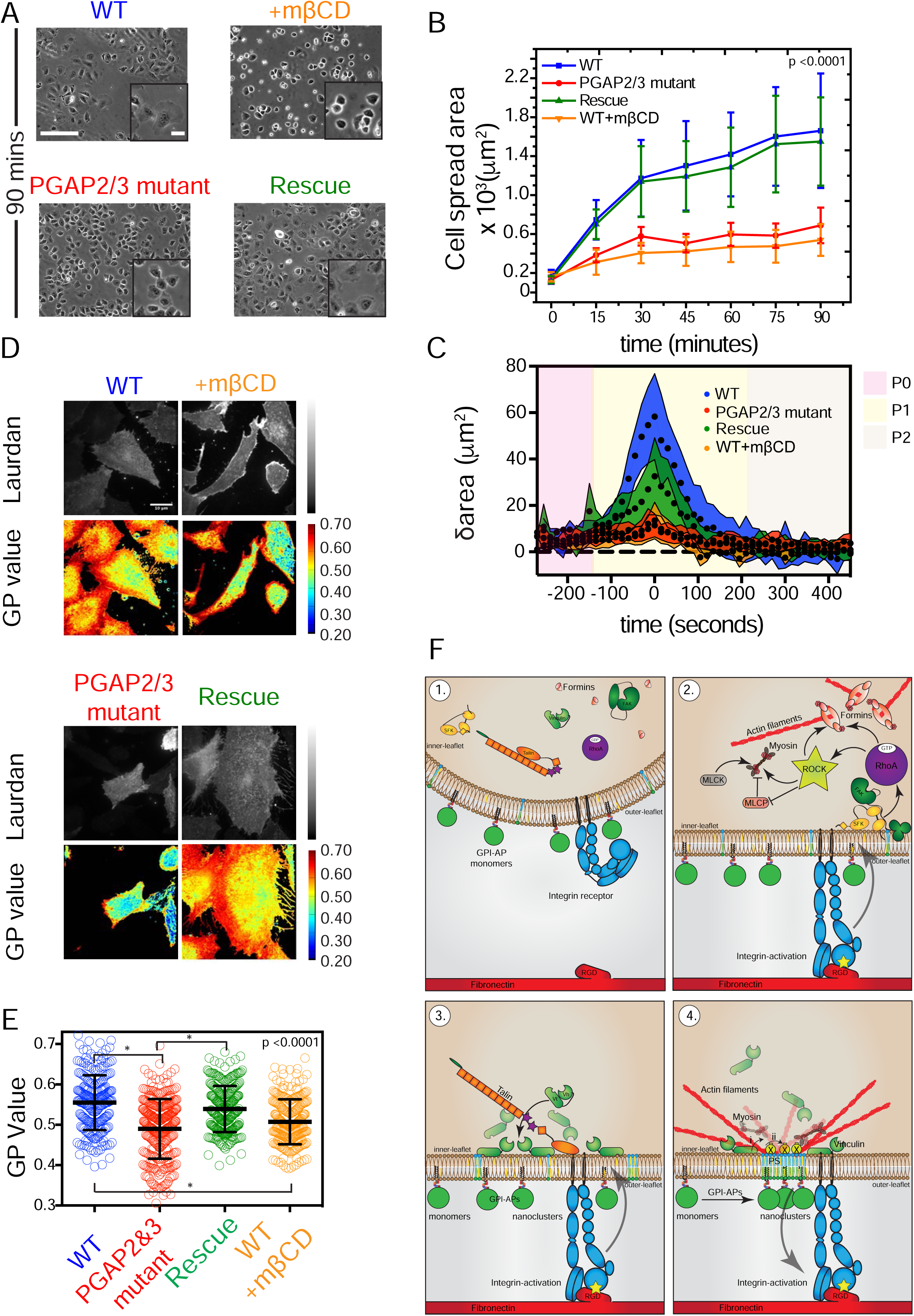
Activity generated GPI-AP nanoclusters are necessary for efficient cell spreading. (A-B) Phase contrast images (A) and corresponding area versus time plot (B) showing the cell spreading dynamics of wild type (blue), PGAP2/3 mutant (red) or PGAP2/3 add back in mutant (Rescue; green) CHO cells or WT cells treated with 10mM mβCD (orange). Each data point is an average of cell spread area of approximately 100 cells per time point. Error bars represent the SD. (C) Plot of change, between two consecutive frames of 15s, in cell spread area (δArea) as a function of cell spreading time on FN for the wild type (blue curve), PGAP2&3 mutant (red curve) and rescue (green curve) in the absence or presence of 10mM mβCD for 30 mins in suspension and during subsequent plating on FN. The data from individual cells was combined by aligning the peak area change (characteristic of the P1 phase) that occurs during individual cell spreading process. Note that the PGAP2&3 mutants and mβCD-treated WT cells lack the rapid increase in cell area characteristic of the P1 phase, consistent with the inability of these cells to make a lamellipodia (Supplementary Movie 3). (D-E) Laurdan total intensity and GP images and (E) Box plot representing the mean generalized polarization (GP) values of WT cells (blue) or PGAP2&3 mutants (red) or PGAP2&3 add-back (Rescue;green) or WT cells treated with 10mM mβCD. The lower values of GP observed in the PGAP2&3 mutants indicate the lack of lo-domains similar to that observed when WT cells were treated with mβCD, a well-characterized perturbant of *lo*-domains. Error bar indicates SD. (E) Model for the integrin signaling triggered generation of nanoclustering of GPI-APs. Integrin ligation triggers a cascade of biochemical signaling events leading to the activation of RhoA via SFK-FAK kinases. This culminates in the activation of the formin class of nucleators that generate dynamic actin filaments along with RhoA-ROCK induced myosin contraction. Vinculin that is activated by talin downstream of integrin activation, either directly (i) or through the activation of additional adapters (ii; X) links the dynamic actin filaments to PS lipids at the inner-leaflet. This in turn couples to the GPI-anchored proteins at the outer-leaflet via a trans-bilayer coupling mechanism. The nanoclusters of GPI-anchored proteins that result from this active mechanism contribute to proper integrin function. Scale Bar 250μm (A) 20μm (inset),10μm (D). Error bar represents SD. See also Figure S7.

Several proteins that either create or reside within *lo*-like regions on the cell membrane have been implicated in the process of cells spreading and migration (Moissoglu et al., 2014; Navarro-Lérida et al., 2012). Recently we have shown that GPI-AP nanoclusters are associated with specific regions of the membrane that have an *lo-* like character (Saha et al, manuscript under preparation). Therefore, we tested if the lack of GPI-AP nanoclusters in the PGAP2/3 mutants could lead to a global disruption of ordered domains. Using the polarity-sensitive membrane dye Laurdan (6-lauryl-2-dimethylamino-napthalene) as a reporter of membrane order (Owen et al., 2012), we found that the PGAP2 and PGAP3 mutant cells have a lower generalized polarization (GP) value (**Figure 7 D-E**), which is consistent with the loss of ordered *lo* domains on the cell surface. The decrease in the GP value was similar to that observed when membrane cholesterol was depleted using mβCD treatment, and is fully restored in the PGAP-2/3 add-back cell line (**Figure 7D-E**). To further confirm that the loss of *lo* domains is due to a specific defect in GPI-AP nanoclustering and not due to global alterations of cholesterol or phospholipid composition of the plasma membrane, we compared the levels of filipin-labeled cholesterol and performed mass-spectrometric measurements on blebs extracted from them. We do not find any significant difference in the levels of either membrane cholesterol or phospholipid profile of the mutant cells (**Figure S7 D-E; Supplementary Table S3**). This suggests that the lack of GPI-AP nanoclustering specifically contributes to the loss of /o-domains at the cell surface and could account for the observable cell spreading defects.

## DISCUSSION

Our results using both chemical and genetic perturbation show how GPI-AP nanoclustering is initiated via a signaling cascade triggered by β1-integrin receptors upon binding to its bonafide ligands: fibronectin (FN) or the fibronectin-derived peptide RGD (see model in **Figure 7F**). Ligand binding results in the activation of the src and FAK kinases. Potentially this step may involve activation of additional molecules including ILK and kindlin-kinases (Calderwood et al., 2013). Regardless, downstream of the kinases are the RhoA GTPases (Ishizaki et al., 1996; Leung et al., 1996), which directly activate formins, necessary for nanoclustering. The nucleator of branched actin filaments, Arp2/3, was not required for this process. Knockdown of both FHOD1 and mDia via RNAi-mediated depletion inhibited nanoclustering of GPI-APs, although the effect of FHOD1 depletion was more drastic. RhoA via ROCK activates FHOD1 through the phosphorylation of C-terminal serine/threonine residues in its DAD region thereby relieving its auto inhibition (Takeya et al., 2008). FHOD1 is also recruited to nascent sites of integrin ligand engagement (Changede et al., 2015; Iskratsch et al., 2013), implicating formins in effecting the nanoclustering of GPI-APs during early stages of integrin mediated signaling. Consistent with this, we found that GPI-AP nanoclusters were also formed on the supported lipid bilayer system, in the vicinity of integrin clusters. Together with previous observations that the integrin LFA1-binding to its ligand ICAM-1 results in a local concentration of GPI-AP nanoclusters, (van Zanten et al., 2009), these results show that integrin signaling generates a localized nanoclustering response.

In parallel, this mechanism requires a way to control myosin function. Indeed, RhoA activates ROCK that regulates myosin light chain kinase (MLCK), and perturbation of ROCK (via Y27632) and MLCK (via ML-7), inhibited actin-based nanoclustering of GPI-APs. However, we do not exclude the possibility of this being a myosin II independent ROCK or MLCK effect. Ectopic activation of RhoA via the agonist CNO3 was sufficient to generate the necessary machinery for creating the GPI-AP nanoclusters, independent of receptor signaling and despite the inhibition of the upstream signaling cascade. Once generated, this actin machinery was also sufficient to cluster transmembrane proteins with actin-binding capacity, exemplified by the model transmembrane protein, FRTM-Ez-AFBD. Transmembrane proteins with actin-binding motifs directly associate with the dynamic acto-myosin machinery, whereas GPI-APs at the outer leaflet, require transbilayer interactions with long acyl-chain containing lipids such as PS at the inner leaflet (Raghupathy et al., 2015). This in turn might require an entirely different mechanism to connect to the actin machinery in the cortex.

Vinculin is known to be activated downstream of integrin signaling through the activation of talin, and has several binding partners such as actin, paxillin and negatively charged lipids like PS and PIP2 (Niggli et al., 1986). The role of these two proteins in supporting GPI-AP nanoclustering was verified by the depletion of talin and vinculin in MEFs, wherein GPI-AP nanoclustering was disrupted. The observation that nanoclusters of TM-ABDs in vin-/- null cells were formed without any alteration implies that the integrin receptor recruits distinct molecular players to facilitate a link between the dynamic actin machinery and membrane lipids. Our results indicate a role for vinculin as a molecular player which may direct actin to the membrane or serve as a linker in connecting the inner leaflet to actin. The failure of the lipid and actin binding mutants to restore GPI-AP nanoclustering support the latter hypothesis. However, vinculin was not found measurably enriched at the membrane outside of focal adhesions when examined at time points where GPI-AP nanoclustering is restored in vin-/- cells (**Figure S6C, D**), supporting the former.

These results provide a molecular mechanism for the control of an active actin-membrane composite, wherein the fluid membrane is inextricably coupled to the cortical actin-substructure beneath (Köster and Mayor, 2016; Rao and Mayor, 2014). The functioning of this composite implies the existence of three types of membrane components: inert, passive and active. While we have previously described inert and passive components and their characteristics (Gowrishankar et al., 2012), here we provide evidence for the functioning of an active element, exemplified by the integrin receptor family.

The relevance of GPI-AP nanoclustering in membrane function has been difficult to probe because of the use of drastic perturbations such as cholesterol removal (Kwik et al., 2003) or alterations in specific phospholipid levels (Lipardi et al., 2000). The identification of a molecular mechanism behind the generation of these nanoclusters, and the key role of integrin signaling in cell spreading provides an opportune physiological context. Pertinently, many of the key components of the nanoclustering molecular machinery identified here such as src, FAK, RhoA, formin, myosin, talin and vinculin, are in the pathway of integrin-mediated signaling, and also have multiple roles in cell physiology. Therefore, the well-documented cell spreading defects and alteration in focal adhesion patterns that are exhibited by perturbations of these players, may be difficult to directly relate to nanoclustering defects. As a consequence, we explored the role of nanoclustering in membrane function by studying defects in integrin-mediated functions in the GPI-anchor remodeling mutants. These mutants lack the ability to make GPI-AP nanoclusters but they support FA-formation as well as integrin-mediated activation. However, they exhibit dramatic defects in cell spreading that are restored upon restoration of the cell’s ability to support nanocluster formation, similar to those observed in vin -/- cells. This implicates a functional role for GPI-AP nanoclustering in integrin-mediated signaling.

Why does signaling via integrin-ligation target the local construction of GPI-AP nanoclusters? An answer to this, is related to the fact that GPI-AP nanoclusters form *lo* nanodomains (Raghupathy et al., 2015) which in turn generate larger meso scale *lo* domains (Saha et al, manuscript in preparation). Here we show that the loss of GPI-AP nanoclustering results in the failure to enhance the overall *lo* characteristics of the membrane observed upon integrin-mediated signaling. Coupled with the observation that large cross-linked patches of GPI-APs accumulate src family kinase members at the inner leaflet (Harder et al., 1998; Stefanová et al., 1991b; Suzuki et al., 2007), these results suggest a function for the *lo*-like GPI-AP nanocluster rich-regions in effecting integrin signaling responses. Lipid modifications such as palmitoylation enable molecules to partition into locally generated *lo* micro-environments. These membrane domains are also likely to be important for the signaling activity of SFK and FAK (Seong et al., 2011). Rac1 is a palmitoylated Rho family GTPase that regulates leading edge protrusion dynamics, and its activity is restricted to *lo* domains (Moissoglu et al., 2014; Navarro-Lerida et al., 2012; del Pozo et al., 2004). Moreover, the GAP activity of the p190RhoGAP is also localized to potentially *lo* domains (Sordella et al., 2003) and its recruitment to *lo*-like regions has been shown to be necessary for the cell spreading process (Arthur and Burridge, 2001), as well as for the localized inhibition of the Rho GTPase and the regulation of FA size. Palmitoylation of the fyn kinase, implicated in rigidity sensing, is also required for the P1-based cell spreading process (Kostic, 2006).Thus, it is likely that the *lo* microenvironment created by the mechanism proposed here, could serve to localize a number of important components of the effector cascade in integrin-based activity to the leading edge where such sensing takes place.

Since the activation of the small GTPase, RhoA, is a pivotal feature downstream of many signaling receptors besides integrins, such as Cadherins, RTKs, GPCRs (Olson and Nordheim, 2010), this will likely culminate in the activation of such an acto-myosin-based mechanism as described here. Vinculin is also a downstream effector of many signaling based systems (Hazan et al., 1997), ensuring that these dynamic acto-myosin filaments also generate GPI-AP nanoclusters. The resultant membrane domains that ensue, will serve as allosteric modulators of the output of the signaling system that generates it (Harding and Hancock, 2008). This will naturally allow the cell to integrate information that is encoded primarily in the composition of its membrane bilayer. In conclusion, our results suggest a generalizable picture of how *lo* nanodomains may be created and deployed in the context of a number of different signaling systems.

## AUTHOR CONTRIBUTIONS

J.M.K., A.A.A., and S.M designed the study. J.M.K, A.A.A set up the EA-TIRFM microscope, performed experiments, and analysed the data; C.P performed the lipid ordering experiments and filipin-labelling experiments; J.M.K also standardized the supported lipid bilayer experiments working in collaboration with M.P.S laboratory. A.A.A also performed the mass spectrometric experiments; T.S.V.Z performed and analysed the FCS measurements and analysed the data involving the cell spreading experiments. J.M.K., A.A.A., and S.M. drafted the manuscript with input from all the authors.

## ACKNOWLEDGMENTS

We thank Cheng-han Yu for help in standardizing the use of the supported-lipid bilayer system; Kabir Husain and Balaji for Matlab codes to analyze the anisotropy and bilayer data; Taroh Kinoshita, Yusuke Maeda, Daniel Rosel, Clare M. Waterman, Lindsey Case, Ana Pasapera for their generous gifts of various reagents (as indicated in the Supplemental Information); Max Planck-NCBS Lipid centre; Bini Ramachandran for mass spectrometry; and H. Krishnamurthy and Manoj Mathew at the Central Imaging and Flow Facility (NCBS). We thank Madan Rao, and Subhasri Ghosh for inspiration and SM lab members for their critical comments on the manuscript. J.M.K. acknowledges pre-doctoral fellowship from NCBS-TIFR. A.A.A acknowledges N-PDF fellowship from DST-SERB (Government of India). T.S.V.Z. acknowledges EMBO fellowship (ALTF 1519-2013) and NCBS fellowship. S.M. acknowledges JC Bose Fellowship from DST (Government of India), a grant from HFSP RGP0027/2012 and Wellcome Trust-DBT Alliance Margadarshi fellowship.

## STAR METHODS

Detailed experimental conditions are provided in the Extended Experimental Procedures in the Supplemental Information.

### Plasmids, Cell Lines, Antibodies and Other Reagents

CHO cells stably expressing EGFP-GPI, Human U2OS cells stably expressing mEGFP-GPI, FR-TM-Ez-AFBD and FR-TM-Ez-AFBD* (RA mutant) cell line, vinculin and Talin 1 deficient mouse embryonic fibroblasts (MEF) and PGAP2/3 double mutant cell lines (of CHO origin) stably expressing CD59 and DAF was maintained in culture media as indicated in the extended experimental procedures with the appropriate antibiotics. The constructs used in the study were procured from various sources as indicated in the extended experimental procedures.

#### GPI analogue incorporation

GPI analogues are incorporated into cell membranes by γ-CD method as described (Koivusalo et al., 2007; Riya Ragupathy doctoral thesis (http://hdl.handle.net/10603/77067).

#### Preparation of RGD functionalized Supported Lipid Bilayers (SLBs)

Supported lipid bilayers functionalized with cRGD was prepared based on the published protocol (Yu et al., 2011).

#### FCS

Fluorescence correlation spectroscopy (FCS) measurements were performed as described previously (Gowrishankar et al., 2012).

### Mass Spectrometry experiments

Briefly, Cells were treated with cytochalasin D in serum free media followed by centrifugation; the collected pellet (membrane blebs) was subjected to lipid extraction using chloroform/methanol solvent system (Raghupathy et al., 2015). The lipid extract thus obtained was subjected to LC/MS by performing mass spectrometric analysis on Q Exactive instrument (Thermo Fisher scientific) (Ramachandran et al, Manuscript in preparation)

#### Anisotropy measurements

Steady state Emission anisotropy-total internal reflection fluorescence microscopy (EA-TIRFM) (Swaminathan et al., 2017). Homo-FRET based anisotropy measurements were carried out on a NikonTE2000 microscope with polarized laser excitation and fitted with an 100x 1.49 NA TIRF objective with a dual camera imaging arrangement as described earlier (Ghosh et al., 2012). Confocal based anisotropy measurements were acquired on a custom-designed NikonTiE microscope coupled to a Yokogawa CSU-22 spinning disc unit (Yokogawa) as described (Ghosh et al., 2012).

#### Cell spreading assay

Briefly, serum-starved cells were de-adhered and plated on pre-cleaned, pre-coated tissue culture dishes under serum-free conditions. After 5 mins of initial cell adherence, the unbound cells were washed off and the dishes were transferred back to 37°C/5% CO_2_ incubator. The cells were imaged in phase contrast with a 20X objective at each of the indicated time points. For quantification of cell-spread area, the cells were marked manually and the mean cell area was extracted using the ROI manager tool in Fiji.

#### Statistical Analysis

Unless otherwise indicated each experimental condition was performed as technical replicates (with at least two dishes per experiment) and data was quantified from >200 (~0.2×0.2μm) region of interest (ROI) taken from 20-30 cells for the anisotropy experiments. Each experiment that has been reported here was performed at least twice with similar results.

Statistical analysis was performed using Matlab (‘ranksum’) functions to determine significance using the Mann-Whitney U test. This is a nonparametric test that assumes unpaired samples that does not necessarily follow normal distributions. Significance levels are indicated as * (p ≤ 0.0001) or n.s (p > 0.0001) where applicable. For details of the sample size and p-values associated with each experimental condition, please refer to **Table S4** and **Table S5** in Supplemental Experimental Procedures.

## SUPPLEMENTARY FIGURE LEGENDS

**Figure S1:**
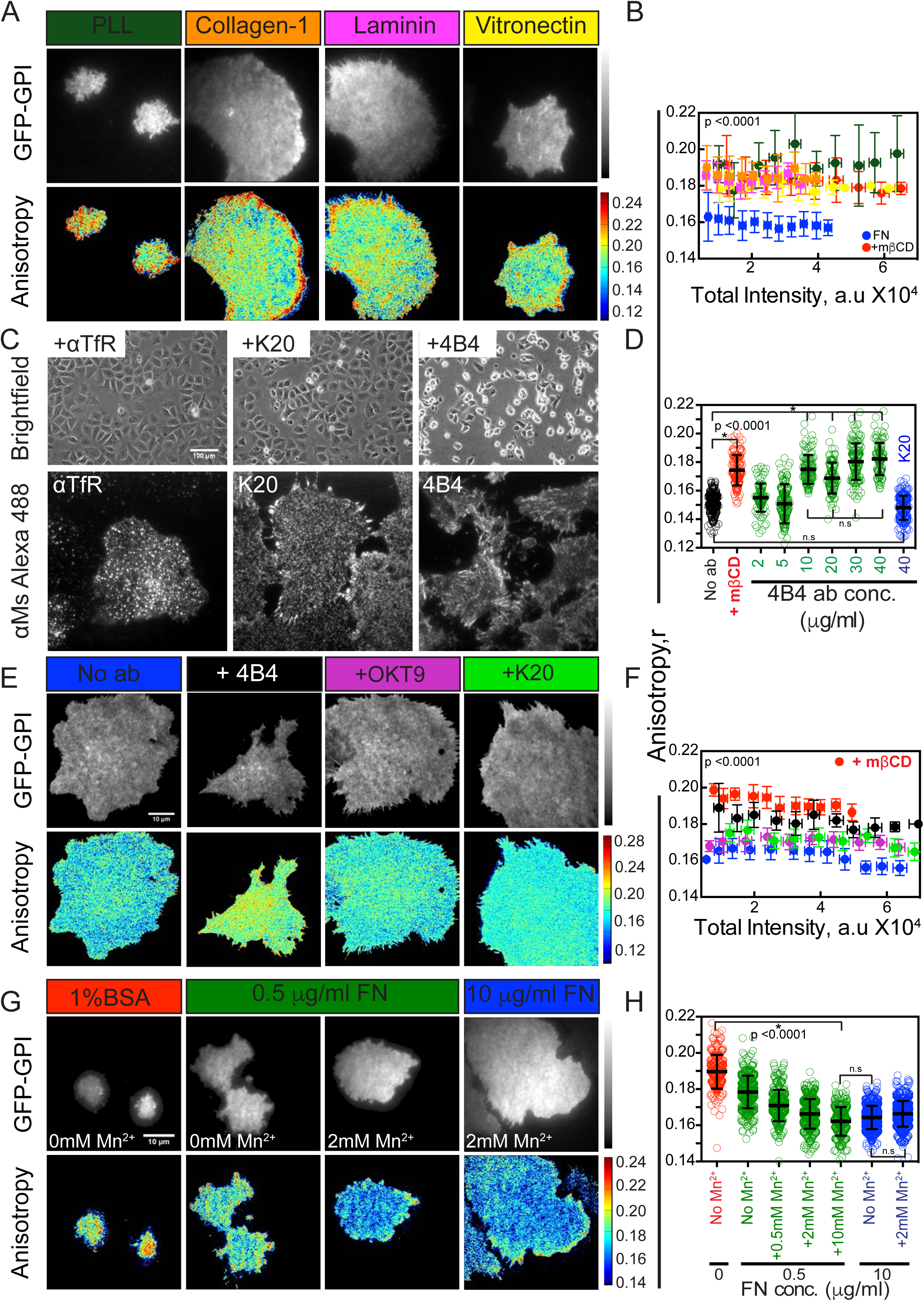
Increase in GPI-AP nanoclustering is dependent on the presence of RGD ligand and occurs in a β-1 integrin dependent manner, Related to Figure 1. (A-B) Intensity and steady state anisotropy images (A) and intensity versus anisotropy plots (B) of GFP-GPI expressing human U2OS cells plated on glass coverslips coated either with 0.01% Poly-L-lysine (PLL; green) or Collagen-1 (orange) or 20μg/ml Laminin or 10μg/ml Vitronectin (yellow) or on 10μg/ml FN (FN; blue) and on FN and treated with 10mM mβCD (mβCD; red) and imaged on EA-TIRFM. Scale bar 10 μm. Note that the fluorescence anisotropy of GFP-GPI in cells plated on FN is highly depolarized compared to cells plated on integrin-inert substrate PLL or on substrates that activates other classes of integrins. (C-D) Treatment of cells with the β1 integrin function antibody alone results in defective cell spreading response (top panel, Brightfield image), implicating β1 integrin as the primary integrin utilized by U2OS cells to spread on FN. These antibodies localize to focal adhesions (C) when probed with appropriate fluorescent secondary antibodies in an immunofluorescence assay (bottom panel, αMs Alexa 488). (D) Box plot depicting mean anisotropy of GFP-GPI in U2OS cells plated on 10μg/ml FN (No ab; black) or pre-treated in suspension with increasing amounts of the β1 integrin function blocking antibody 4B4 (2-40μg/ml; green) or with 40 μg/ml of a neutral (non-function perturbing) β1 integrin antibody K20 (blue) and subsequently plated on FN in the presence of the respective concentrations of the antibodies and imaged on EA-TIRFM. Scale bar 10 μm. Note that while blocking the function of β1 integrin disrupts the increase in nanoclustering of GPI-APs seen on fibronectin, blocking the αV integrins does not result in the loss of GPI-AP nanolcusters (Data not shown). (E-F) Intensity and steady state anisotropy images (E) and intensity versus anisotropy plots (F) of GFP-GPI expressing U2OS cells plated on FN (No ab, blue) or pre-treated in suspension with 40 μg/ml of a neutral (non-function perturbing) β1 integrin antibody K20 (Green) or 40 μg/ml of an antibody against the transferrin receptor (OKT9, magenta) and subsequently plated on 10μg/ml FN in the presence (red) or absence of 10mM mμCD and imaged on EA-TIRFM. Scale bar 10μm. (G-H) Intensity and steady state anisotropy images (G) and (H) Box plot depicting the mean anisotropy of GFP-GPI in U2OS cells plated on glass blocked with 1%BSA (red) without Mn^2+^ or on 0.5μg/ml FN (green) with or without 2mM Mn^2+^or on 10μg/ml FN (green) with 2mM Mn^2+^ (blue) imaged on EA-TIRFM. Scale bar 10 μm. Note that shifting the equilibrium towards ligand-engaged integrin, either by increasing FN density or by the activation of integrin by Mn^2+^ promotes the generation of GPI-AP nanoclusters. Error bars represent SD.

**Figure S2:**
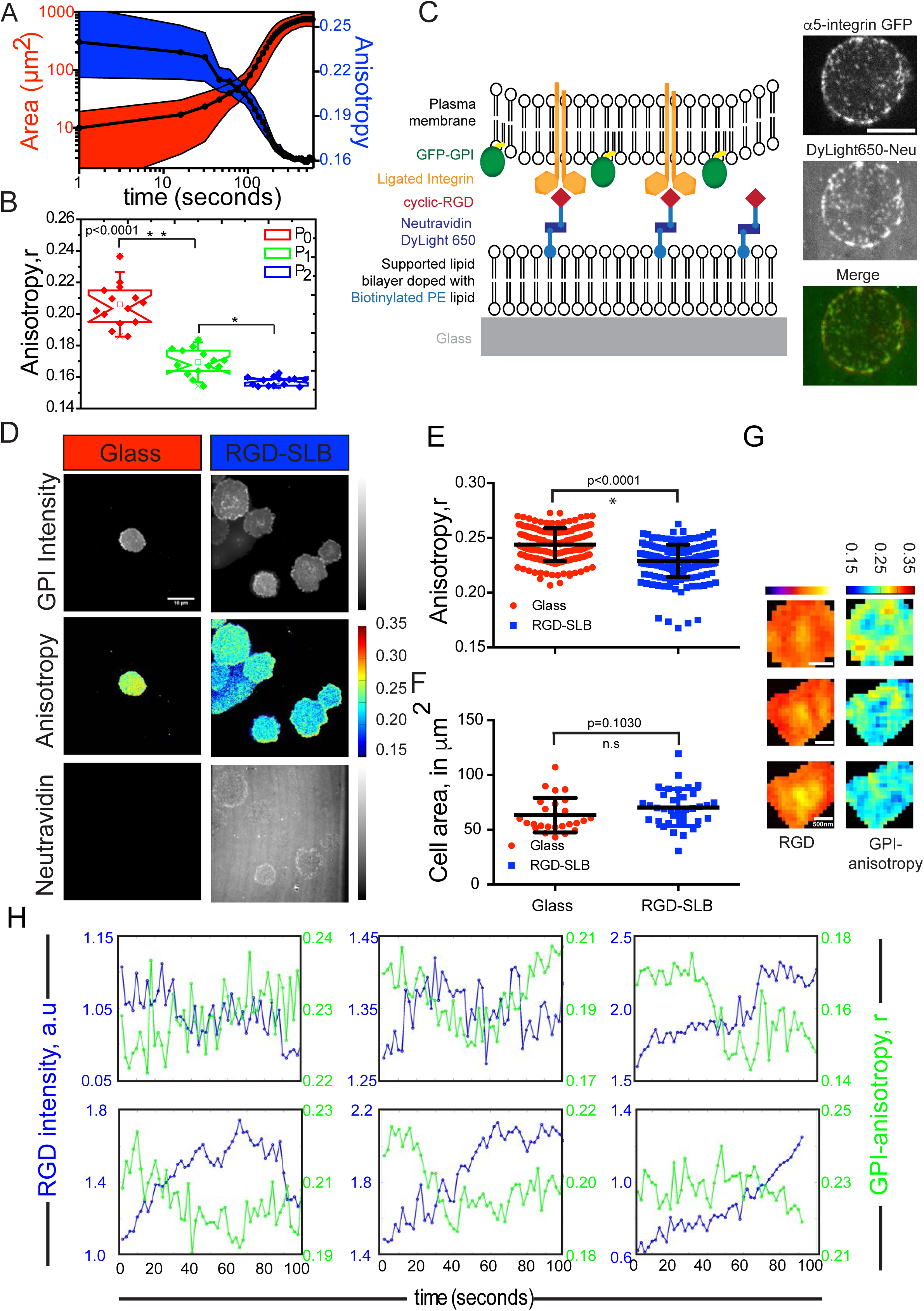
Activation of RGD binding integrins leads to enhanced nanoclustering of GPI-APs in its local vicinity, Related to Figure 2. (A) Plot of cell spread area (red curve) with corresponding change in the anisotropy of GFP-GPI (blue curve) as a function of spreading time (log time-X axis) of CHO cells expressing GFP-GPI taken using EA-TIRFM. (B) Notch-box plot of the mean anisotropy of cells in the indicated phases of cell spreading quantified by drawing ROIs in the intensity kymographs and extracting the corresponding values from the anisotropy kymographs. In the box plots, box includes the median and the 1^st^ quartile to 3^rd^ quartile. Whisker extends 1.5 SD (C) Schematic representation of the RGD-functionalized supported lipid bilayer (SLB) system. Lipids used to prepare the SLB were 1,2-dioleoyl-sn-glycero-3-phosphocholine (DOPC) doped with 1,2-dipalmitoyl-sn-glycero-3-phosphoethanolamine-N-(cap biotinyl) (16:0 Biotinyl Cap PE) (Bottom bilayer). Biotin-cRGD attached to DyLight 650 Neutravidin was used as a linker to facilitate the attachment of GFP-GPI and integrin expressing cells onto the SLB. The DyLight 650 neutravidin signal serves as a marker for the α5(β1) integrin clusters since they co-localize to α5 integrin clusters formed on α5-GFP expressing CHO-B2 cells (right panel; merge). Scale bar 5 μm. (D) Intensity and anisotropy images and (E, F) graphs representing the anisotropy (E) and cell spread area (F) of GFP-GPI expressing cells plated on RGD functionalized supported lipid bilayers (blue) or glass (red) show that the anisotropy of cells plated on cRGD functionalized SLBs were lower than those of cells plated on plain glass under comparable cell spread area. Error bars are SD. (G) Representative montages of ROIs demonstrating correlations between RGD cluster intensity (more ‘yellow’ pixels denote higher cluster intensity) with corresponding GFP-GPI anisotropy maps (more ‘blue’ pixels represent regions with more GPI nanoclusters) taken from cells expressing GFP-GPI re-plated on RGD functionalized SLBs. Scale bar 500 nm. (H) Representative plots that demonstrate correlations between the RGD-cluster intensity (blue curve) and GFP-GPI anisotropy (green curve) and taken over time from ROIs sampled from 30 cells (20-30 clusters per cell) plated on RGD-functionalized fluid-SLBs. Note that the increase (or decrease) in RGD cluster intensity coincides with a decrease (or increase) in GPI-AP anisotropy indicating a local increase (or decrease) in GPI-AP nanoclusters. Error bar represent SD.

**Figure S3:**
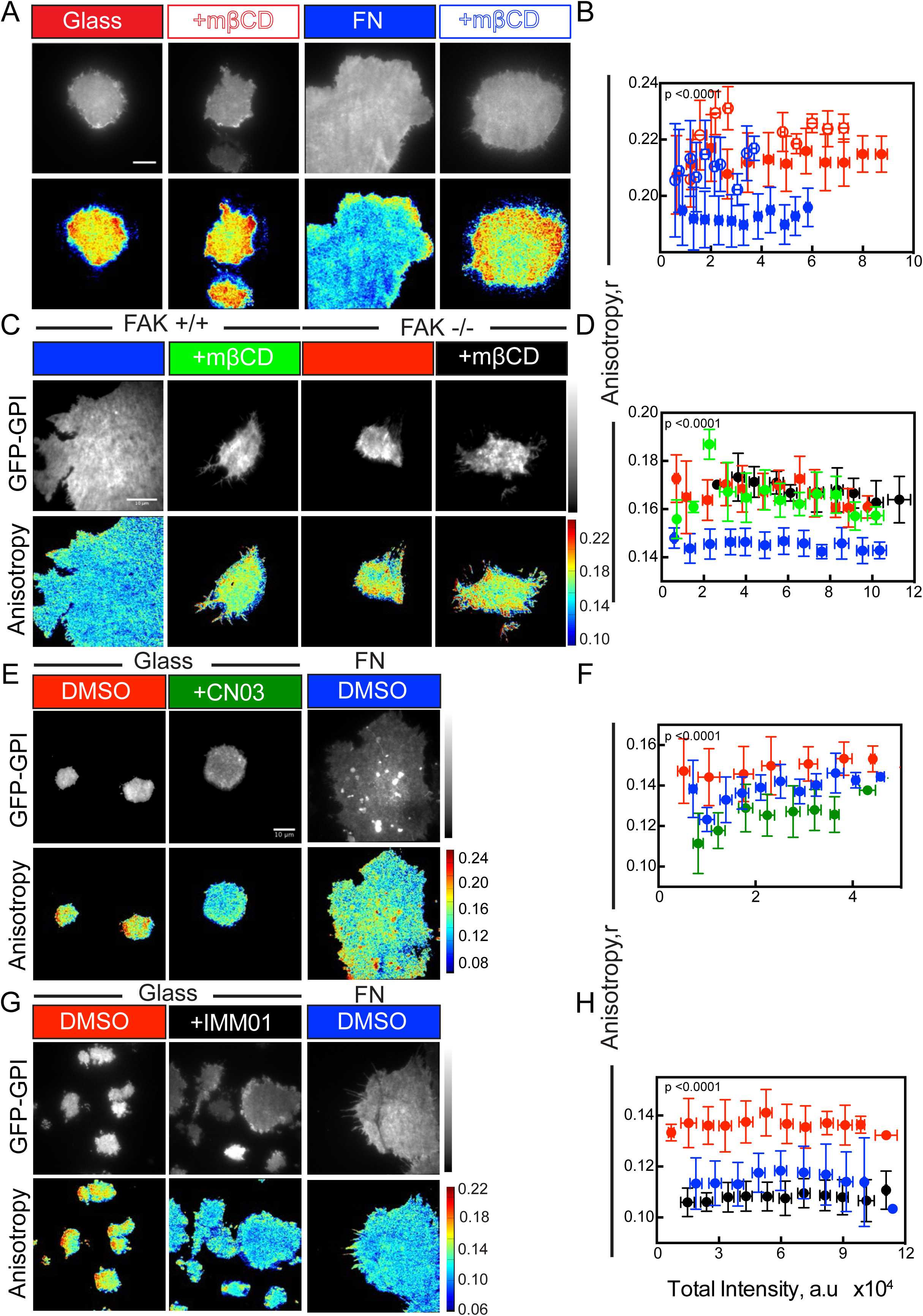
Effect of perturbations of downstream targets of integrin signaling on GPI-AP nanoclustering, Related to Figure 3. (A-B) Intensity and steady state anisotropy images (A) and intensity versus anisotropy plots (B) of GFP-GPI expressing CHO cells on glass (red closed circles) or cells on glass and treated with 10mM mβCD (red open circles) or plated on FN (blue closed circles) or plated on FN and subsequently treated with 10mM mβCD (blue open circles. Note that the anisotropy of cells on glass closely resemble those of cells treated with mβCD and therefore either of these conditions can be used to represents cells with a loss of nanolcusters of GPI-APs. (C-D) Intensity and steady-state anisotropy images (C) and intensity versus anisotropy plots (D) of GFP-GPI expressing FAK+/+ (blue or green) or FAK-/- (red or black) mouse embryonic fibroblasts (MEFs) cells plated on 10μg/ml FN in the presence (green or black) or absence (blue or red) of 10mM mβCD imaged on a EA-TIRFM. (E-F) Intensity and steady state anisotropy images (E) and intensity versus anisotropy plots (F) of GFP-GPI expressing CHO cells plated on glass(red) or pretreated and plated on glass in the presence of 10μg/ml RhoA activator (CN03) (green) or plated on FN (blue) and imaged on a TIRF microscope. Note that the fluorescence anisotropy of GFP-GPI in cells treated with the RhoA activator is lower compared to DMSO treated cells on glass and resembles cells plated on FN indicating an increase in GPI-AP nanoclustering independent of FN binding under this condition.(G-H) Intensity and steady state anisotropy images (G) and intensity versus anisotropy plots (H) of GFP-GPI expressing CHO cells plated on glass(red) or pre-treated and plated on glass in the presence of 10μM formin activator (IMM01) (black) or plated on FN (blue) and imaged on EA-TIRFM. Note that the fluorescence anisotropy of GFP-GPI in cells treated with the formin activator is lower compared to DMSO treated cells on glass and resembles cells plated on FN indicating an increase in GPI-AP nanoclustering independent of FN binding under this condition. Error bars indicate SD. Scale bar 10μm.

**Figure S4:**
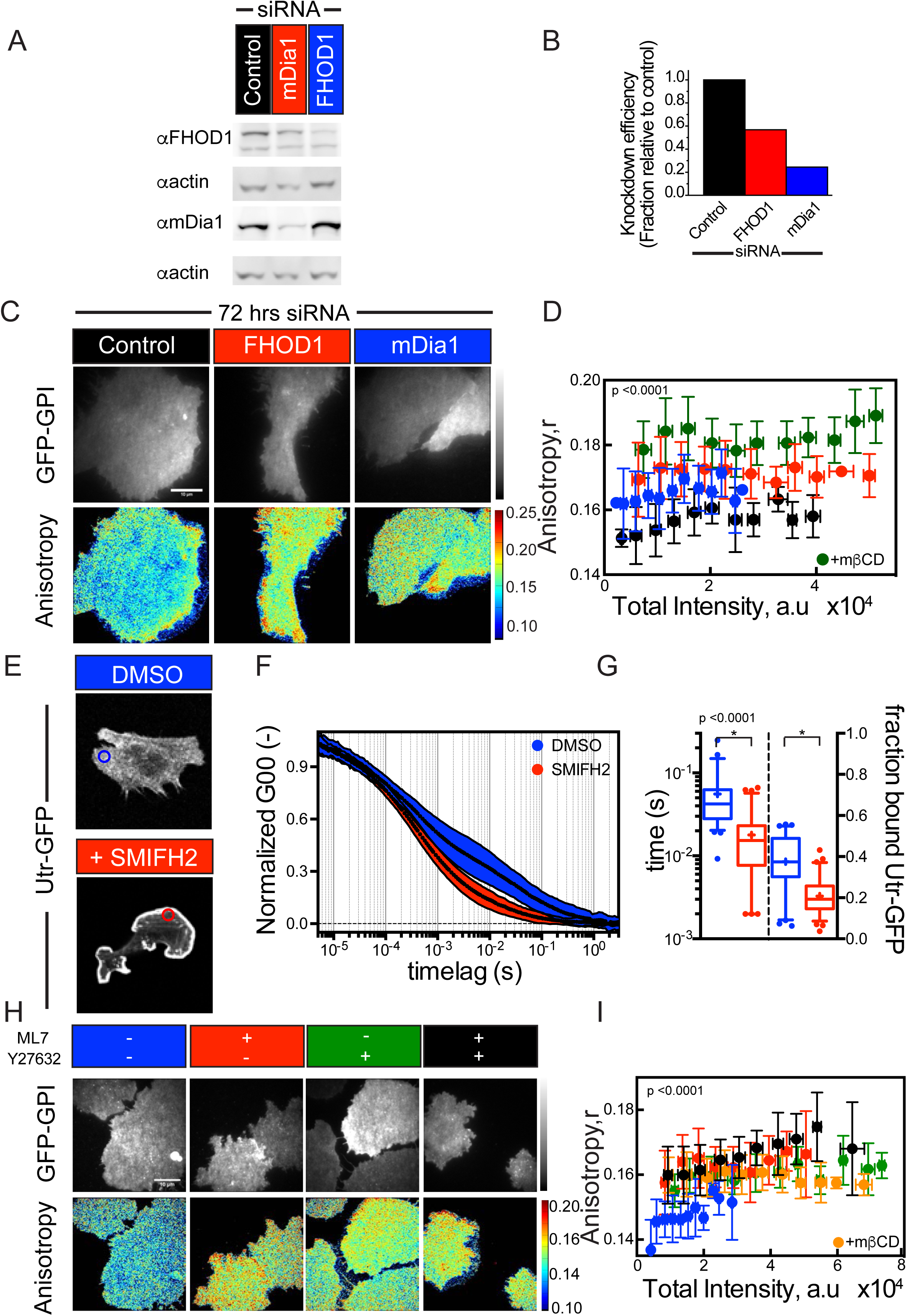
Integrin activation alters cortical acto-myosin activity, Related to Figure 4. (A) Western blot analysis of siRNA medicated knockdown of formins in human U2OS cells transfected for 72 hours with 5nmoles SMART pool of either control scrambled siRNA (black) or siRNA against the formins FHOD1 (red) or mDia1 (DIAPH1; blue) and probed with antibodies against the same. (B) Quantification of the blots indicate a knockdown of ~45% FHOD1 protein levels and ~80% in the case of mDia. The data is normalized first to the corresponding intensities of β-actin in each well (loading control) and then to the levels of the protein in the control siRNA well. (C-D) Intensity and steady-state anisotropy images (C) and intensity versus anisotropy plot (D) of U2OS cells expressing GFP-GPI plated on 10μg/ml FN after treatment with 5nmol of either control siRNA (blue) or FHOD1 siRNA (red) or mDia1 siRNA (green) for 72 hours and imaged in EA-TIRFM. Scale bar 10μm. Error bars indicate SD. (E-G) Confocal images (E) and Average (circles) and standard deviation (shaded) autocorrelation decays (F) of GFP-tagged Utrophin actin filament binding domain (GFP-Utr) in cells plated on FN (blue) or pre-treated and plated on FN in the presence of 10μM formin inhibitor SMIFH2 (red). FCS data was collected from regions in the cell periphery devoid of stable actin filaments as indicated (blue or red circles). Note the loss of the slow diffusion timescales corresponding to 10ms or longer for cells treated with the formin inhibitor. (G) Timescale and fraction of the slow moving population associated with moving actin filaments. (H-I) Intensity and steady-state images (H) and intensity versus anisotropy plot (I) of CHO cells stably expressing GFP-GPI plated on 10μg/ml FN after treatment with DMSO (control; blue) or 20μM MLCK inhibitor ML7 (yellow) or 20μM ROCK inhibitor Y-27632 (green) or 20μM of both ROCK and MLCK inhibitors (ML+Y;black) or mβCD (orange) taken on EA-TIRFM. Scale bar 10 μm. Error bars indicate SD.

**Figure S5:**
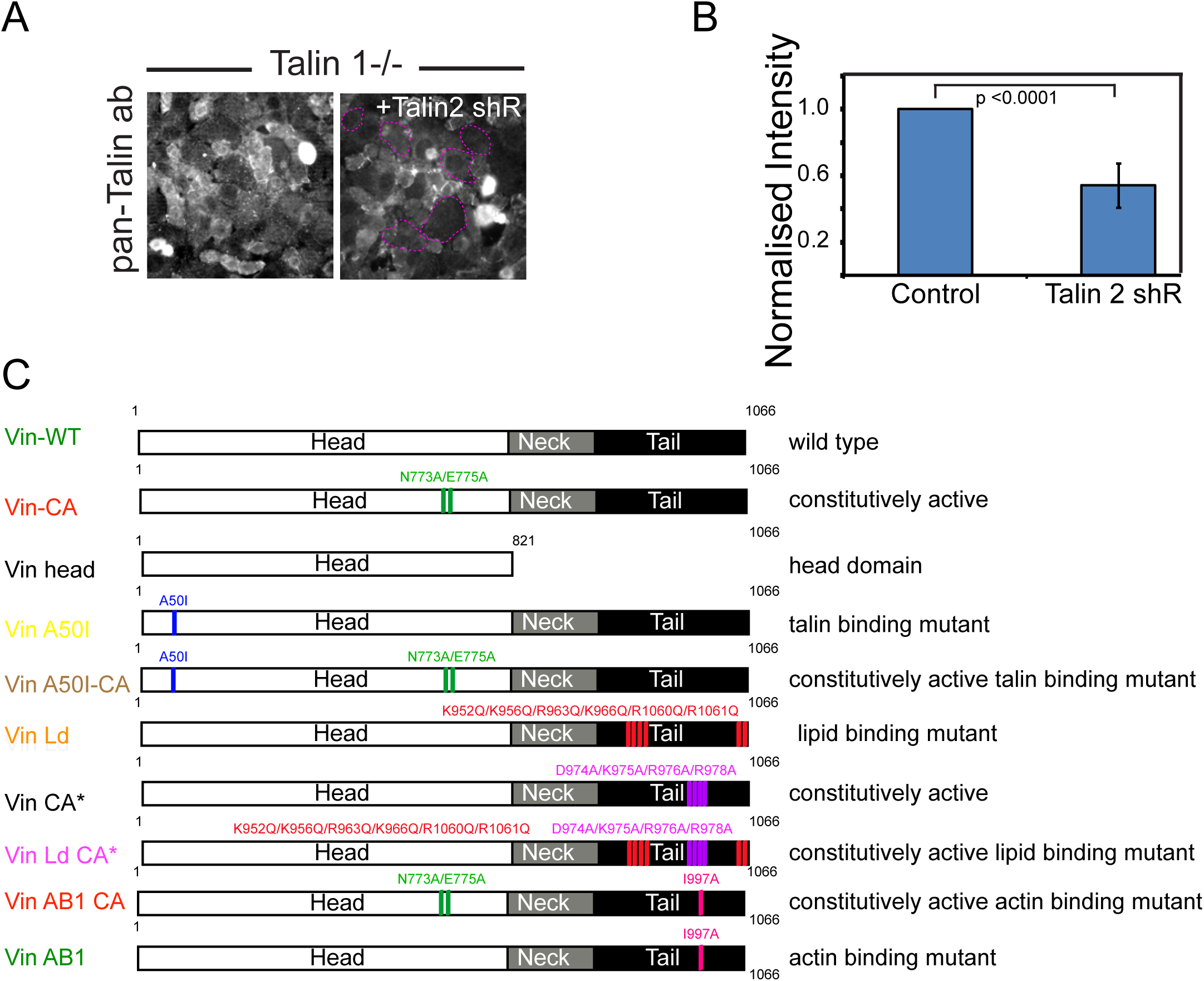
Characterization of vinculin deficient and talin 1 deficient MEFs and schematic of vinculin mutations used in this study, Related to Figure 5. (A-B) Intensity images (A) and corresponding histogram (B) quantifying endogenous talin levels in talin 1-/- cell line in the presence or absence of GFP-tagged talin 2 shRNA using a pan talin antibody. The data shows a significant decrease in talin levels in cells treated with talin 2 shRNA (C) Schematic depicting the vinculin variants used in this study, and the typical characteristics of each molecule. Error bars represent SD.

**Figure S6:**
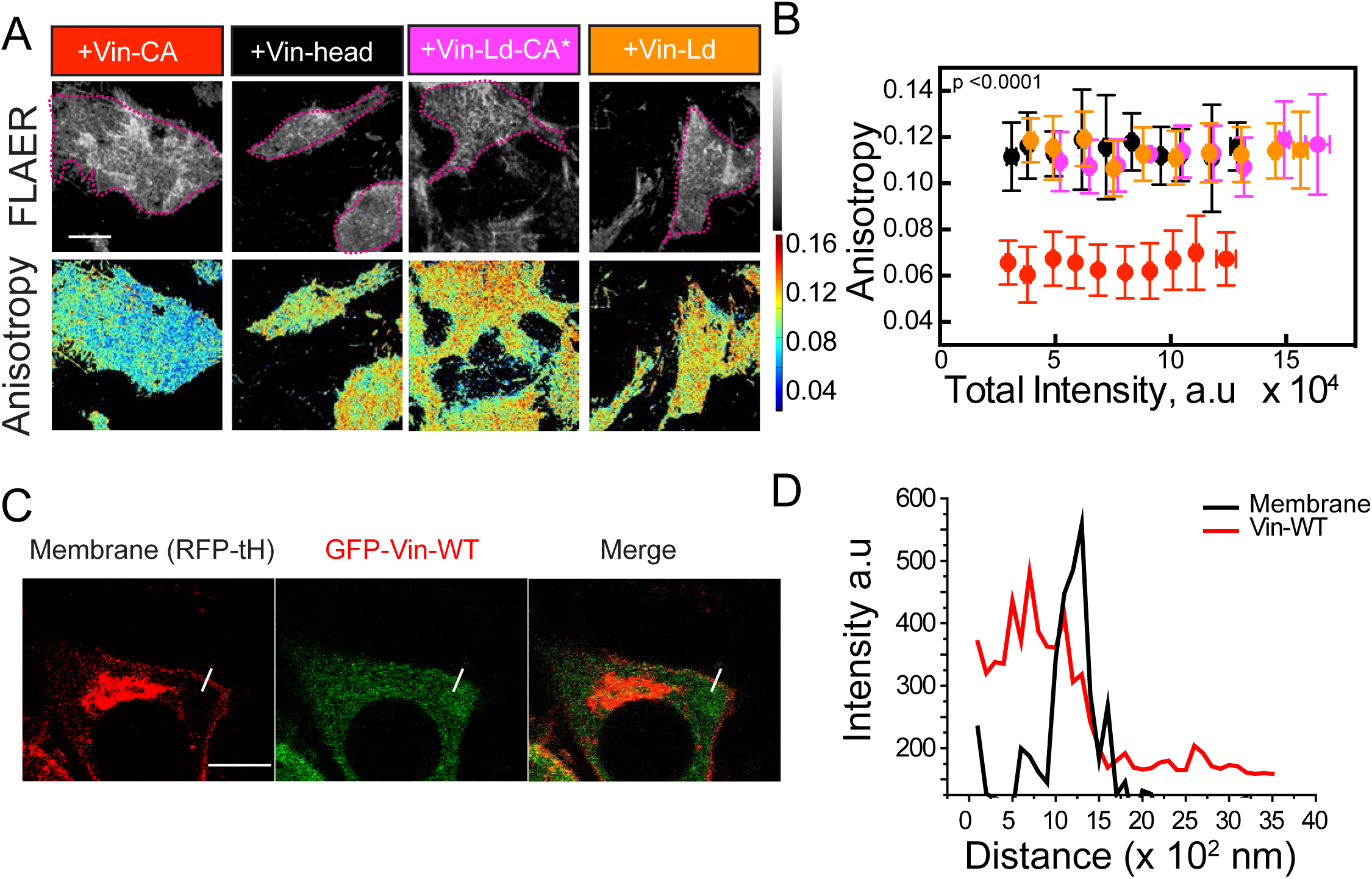
Characterization of the effects of vinculin mutants on GPI-AP nanoclustering, Related to Figure 6. (A-B) Intensity and anisotropy images (A) and intensity versus anisotropy plot (B) of vinculin deficient (vin-/-; MEFs transfected with the indicated vinculin variants and labeled with Alexa-568-FLAER 12-16 hours post transfection and imaged in EA-TIRFM after replating the cells on FN coated glass bottom dishes. Transfected cells are marked by dotted magenta lines. (C-D) vin-/- MEFs were transfected with GFP-tagged Vinculin and co-transfected with a membrane marker (RFP-tHRas) and imaged on a confocal microscope (C). Plot (D) shows the line intensity profiles of Vin-WT and RFP-tHRas suggesting the absence of Vin-WT at the plasma membrane post 12-16 hours of transfection. White line in (C) depicts the region of line scan measurement. Scale bar 10 (A), 5 (C)μm. Error bar represent SD.

**Figure S7:**
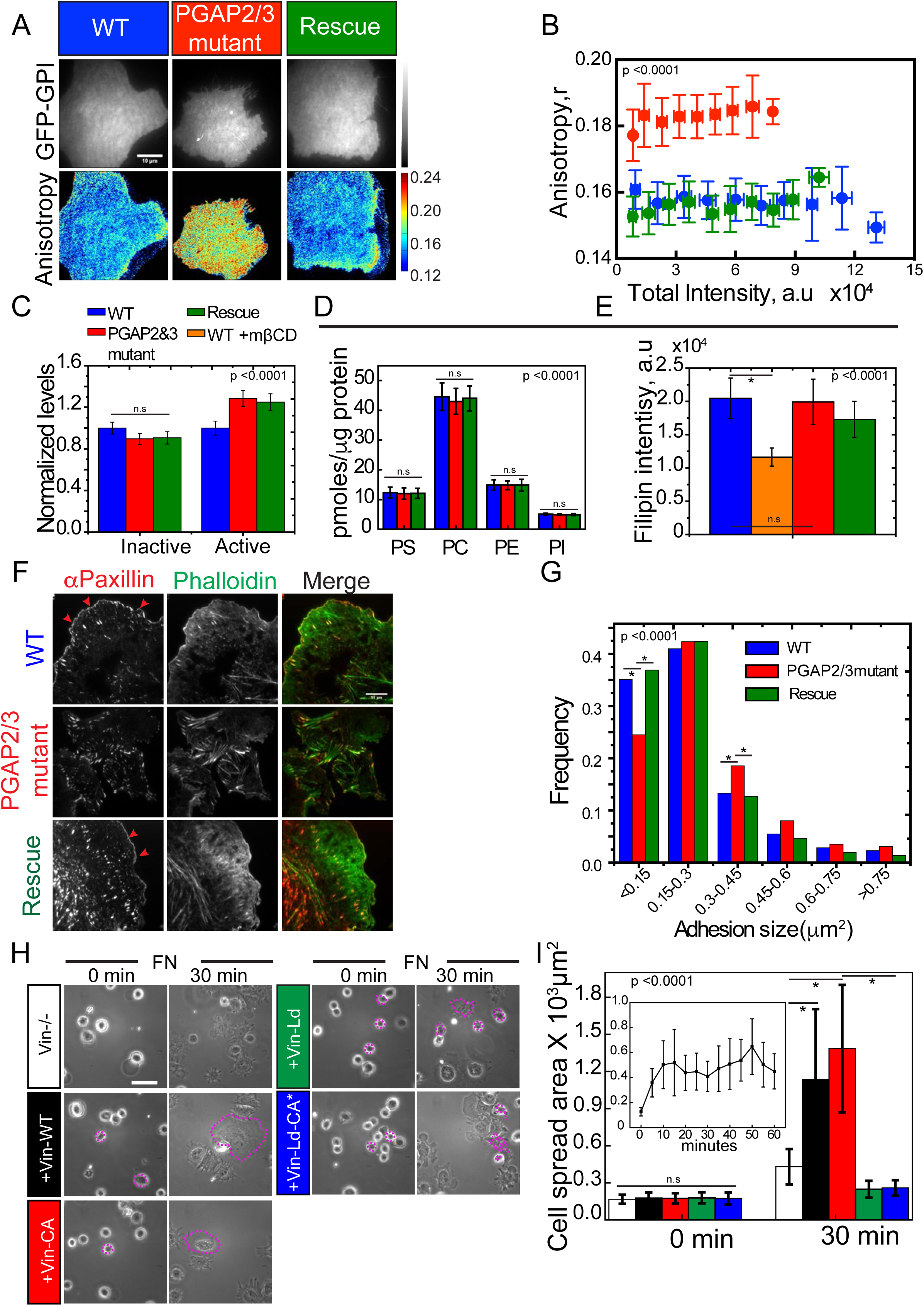
Functional significance of GPI-AP nanocluster formation, Related to Figure 7. (A-B) Intensity and steady-state anisotropy images (A) and intensity versus anisotropy plot (B) of GFP-GPI expressed in wild type (WT; blue), PGAP2/3 double mutant (red) and PGAP2/3 add back (Rescue; green) re-plated on 10μg/ml FN. Scale bar 10μm.Note that an increase in anisotropy in mutant cells was observed corresponding to a loss of nanoclustering of GFP-GPIs. (C) Quantification of the relative amount of active β-1 integrins (marked by HUTS4 antibody) and inactive β-1 integrins (marked by 4B4 antibody) normalized to the levels of neutral antibody (K20) of WT cells (blue bar), PGAP2&3 mutants (red bar), rescue cells (green bar). Data is also additionally normalized to the levels in the WT scenario. (D-E) MS/MS Mass spectrometric analysis (D) and filipin-staining of WT cells (blue bar) or PGAP2&3 mutants (red bar) or rescue (green bar) or WT cells treated with 10mM mβCD (control in E) quantifying the levels of the indicated lipid species (in D) and free cholesterol (in E). Data represents the mean +/- SD. (F) TIRFM images of WT (blue), PGAP2&3 double mutant (red) and Rescue cells (green), de-adhered and allowed to spread on FN coated dishes for 60 mins and subsequently fixed, permeabilised and stained for paxillin (Left panel) to mark focal adhesions, or labeled with Phalloidin to mark actin filaments (Middle panel) or a merge of both (Right panel. Scale bar 10μm. (G) Frequency histogram of the binned focal adhesion sizes (marked by paxillin) of WT (Blue bars), PGAP2/3 double mutant (Red bar) and Rescue line (Green bars) Note that the mutant cells have larger adhesions and lack the smaller nascent adhesions that are usually found at the cell periphery/lamellipodia (Red arrow heads in F). (H-I) Phase contrast images (H) and histogram (I) quantifying the extent of spreading of vin-/- MEFs or vin-/- MEFs transiently transfected with the indicated Vin constructs. Cell area was quantified at 0 and 30 mins after seeding on 10μg/ml FN-coated glass-bottom dishes. Inset depicts the cell spreading versus time profile of vin-/- cells on FN. The transfected cells are marked by a dotted magenta line. Scale bar 50 μm. Error bars represent SD.

## Supplementary Movie Legends

**Supplementary Movie 1:** Cell spreading dynamics of GFP-GPI expressing CHO cells on 10μg/ml FN; Imaged after every 15 seconds in 100X EA-TIRFM mode at 37°C. Left panel: Total Intensity image; Right Panel: GFP-GPI anisotropy image with the corresponding LUT bar. Scale bar 10μm. Notice that the cells initially blebs and rapidly acquire GPI-AP nanoclusters (blue pixels) before the cell extends out a prominent lamellipodia and begins spreading rapidly.

**Supplementary Movie 2:** 20X Phase contrast time series images of WT or PGAP2&3 double mutant CHO cells spreading on FN. Notice that the PGAP2&3 double mutants spread slowly and extend by producing blebs (and lack a prominent lamellipodia). Scale bar 50μm.

**Supplementary Movie 3:** Cell spreading dynamics of GFP-GPI (Cell membrane marker) expressing WT (Left Panel), PGAP2&3 mutant (Middle Panel) or Rescue (Right Panel) CHO cells spreading on 10μg/ml FN;imaged every 15 seconds in 100X TIRF at 37°C. Notice that the PGAP2&3 mutant cells exhibit defects in cell spreading due to their inability to produce a prominent lamellipodia. Scale bar 10μm.

